# The Alzheimer’s disease gene *SORL1* regulates lysosome function in human microglia

**DOI:** 10.1101/2024.06.25.600648

**Authors:** Swati Mishra, Nader Morshed, Sonia Sindhu, Chizuru Kinoshita, Beth Stevens, Suman Jayadev, Jessica E. Young

## Abstract

The *SORL1* gene encodes the sortilin related receptor protein SORLA, a sorting receptor that regulates endo-lysosomal trafficking of various substrates. Loss of function variants in *SORL1* are causative for Alzheimer’s disease (AD) and decreased expression of SORLA has been repeatedly observed in human AD brains. *SORL1* is highly expressed in the central nervous system, including in microglia, the tissue resident immune cells of the brain. Loss of SORLA leads to enlarged lysosomes in hiPSC-derived microglia like cells (hMGLs). However, how SORLA deficiency contributes to lysosomal dysfunction in microglia and how this contributes to AD pathogenesis is not known. In this study, we show that loss of SORLA results in decreased lysosomal degradation and lysosomal enzyme activity due to altered trafficking of lysosomal enzymes in hMGLs. Phagocytic uptake of fibrillar amyloid beta 1-42 and synaptosomes is increased in SORLA deficient hMGLs, but due to reduced lysosomal degradation, these substrates aberrantly accumulate in lysosomes. An alternative mechanism of lysosome clearance, lysosomal exocytosis, is also impaired in *SORL1* deficient microglia, which may contribute to an altered immune response. Overall, these data suggest that SORLA has an important role in proper trafficking of lysosomal hydrolases in hMGLs, which is critical for microglial function. This further substantiates the microglial endo-lysosomal network as a potential novel pathway through which *SORL1* may increase AD risk and contribute to development of AD. Additionally, our findings may inform development of novel lysosome and microglia associated drug targets for AD.

## 1 INTRODUCTION

Treatments for Alzheimer’s disease (AD), a neurodegenerative disorder and the most common cause of dementia (“2023 Alzheimer’s disease facts and figures,” 2023), are few and do not alter the course of disease progression by more than several months. Hallmarks of AD include extracellular Amyloid-beta (Aβ) accumulation in senile plaques, intracellular aggregation of tau protein in neurofibrillary tangles, neurodegeneration and neuroinflammation. Microglia, tissue-resident phagocytes of the brain, are key effectors of pathogenic protein clearance and maintenance of neuronal health and brain homeostasis in general, positioning this brain cell type at the center of cellular responses in AD pathogenesis and making it an attractive target for new AD therapies. Identification of AD risk genes highly expressed in microglia by genome-wide association studies (GWAS) further supports the idea that dysfunctional microglia may contribute to onset and/or progression of AD(Maninger et al., 2024; McQuade & Blurton-Jones, 2019). Collective evidence from murine models and post-mortem human AD brain tissue has revealed that microglia migrate to and cluster around Aβ plaques(Akiyama et al., 1999; Bolmont et al., 2008; Itagaki et al., 1989; Meyer-Luehmann et al., 2008; Stalder et al., 1999) but are unable to eliminate these toxic deposits from the brain. One mechanism that may cause this defect is lysosome dysfunction in microglia. Abnormal lysosomes in microglia can negatively alter phagocytic and inflammatory responses (Iyer et al., 2022; Lancaster et al., 2021; Joseph D. Quick et al., 2023; Ren et al., 2022), functions known to cause neuroinflammation in AD, substantiating the need to investigate microglia-specific lysosomal pathways as therapeutic targets. Our previous work has shown that loss of the intracellular sorting receptor and potent AD risk gene, Sortilin-related receptor-1 (*SORL1*), that encodes the multidomain protein SORLA, results in enlarged lysosomes in hiPSC-derived microglia (Mishra et al., 2024).

SORLA is a hybrid receptor that belongs to both the LDLR and the VPS10p receptor family. It interacts with the multi-protein assembly retromer and shuttles proteins between TGN, endosomes and cell surface (Fjorback et al., 2012; Jacobsen et al., 2001; Jacobsen et al., 1996; Willnow & Andersen, 2013; Yamazaki et al., 1996). In particular, retromer and its receptors, SORLA and the cation-independent mannose-6 phosphate receptor (CIMPR), function to traffic mature lysosomal hydrolases from the TGN via endosomes to lysosomes (Cui et al., 2019; Hasanagic et al., 2015; Nielsen et al., 2007). Genetic, epidemiological and functional studies have now cumulatively and unequivocally established *SORL1* as a gene that increases risk for AD, highlighting aberrant endo- lysosomal trafficking as an important pathway in AD pathogenesis (Holstege et al., 2017; Holstege et al., 2023; Kunkle et al., 2017; Lambert et al., 2013; Rogaeva et al., 2007; Scheltens et al., 2021). Loss of SORLA expression has been observed in AD brains in the early stages of the disorder (Dodson et al., 2006; Scherzer et al., 2004; Thonberg et al., 2017). Studies investigating the role of SORLA in AD pathogenesis have extensively focused on neurons and revealed that loss of SORLA causes endo-lysosomal defects in neurons, specifically in endosomal recycling, autophagy and lysosome function, ultimately resulting in increased secretion of toxic Aβ species and impaired synaptic activity in neurons(Andersen et al., 2022; Hung et al., 2021; Knupp et al., 2020; Lee et al., 2023; Mishra et al., 2023; Mishra et al., 2022). While *SORL1* mRNA is highly expressed in microglia, even more so than neurons and astrocytes(Lee et al., 2023; Olah et al., 2018; Zhang et al., 2016), the impact of loss of SORLA function specifically in microglia is not well-studied, rendering our understanding of the role of SORLA in AD risk incomplete at best. Understanding cell type specific functions of AD risk genes, including SORLA, is imperative in understanding the etiology and pathogenesis of the disease, and to develop rational therapeutic approaches for AD.

In this study, we investigated how SORLA deficiency affects lysosomal function in hiPSC-derived microglia like cells (hMGLs). We used quantitative flow cytometry, immunocytochemistry, and functional enzymatic and cellular assays to reveal that *SORL1* KO hMGLs show decreased lysosomal degradation and reduced lysosomal enzymatic activity, due to impaired trafficking of lysosomal enzymes from the trans-Golgi network (TGN) to lysosomes. We show that the main retromer cargo that functions with SORL1 to traffic lysosomal enzymes, CIMPR, has reduced expression in SORL1 deficient hMGLs and reduced co-localization with enzymes in lysosomes. This impairment in enzyme trafficking leads to inefficient lysosomal degradation and accumulation of phagocytosed fibrillar Aβ 1-42 and synaptosomes in lysosomes. Impaired degradation leads to lysosomal stress and could prompt cells to rid cells of undigested cargo via other mechanisms, (Domingues et al., 2024; Wang et al., 2018; Zhong et al., 2023). We tested lysosomal exocytosis (LE), a secretory pathway through which microglia release lysosomal enzymes, cargo and inflammatory cytokines into the extracellular environment and found that this process is also altered in *SORL1* KO hMGLs. Overall, our data show that *SORL1* regulates lysosomal homeostasis in human microglia. Loss of normal *SORL1* function leads to mis- trafficking of lysosomal enzymes and impairment of critical lysosomal functions, including substrate degradation and lysosomal exocytosis. This data, along with recent work describing a role of SORLA in sorting neuroimmune receptors (Ovesen et al., 2024), adds to a growing body of work highlighting *SORL1* and related pathways as novel therapeutic targets for AD.

## 2. MATERIALS AND METHODS

### 2.1 Cell lines

*Cell lines generated by CRIPSR/Cas9 gene editing technology* Isogenic cell lines with loss of SORL1 were generated using CRISPR/Cas9 genome editing as described in our previous study (Knupp et al 2020). Briefly, cell lines were generated from our previously published and characterized CV background human induced pluripotent stem cell line (Young et al., 2015). This cell line is male and has a APOE ε3/ε4 genotype (Levy et al., 2007). Genome edited lines were generated using published protocols (Young et al., 2018). Briefly, guide RNAs (gRNAs) to SORL1 were generated using the Zhang Lab CRISPR Design website at MIT (http://zlab.bio/guide-design-resources) and selected to minimize off-target effects. gRNAs were cloned into vector px458 that co-expresses the Cas9 nuclease and GFP, and hiPSCs were electroporated with the plasmid. Electroporated hiPSCs were FACS sorted for GFP, plated in 10 cm plates at a clonal density (∼1x10^4 cells/plate), and allowed to grow for roughly 2 weeks. Colonies were picked into 96 well plates and split into two identical sets. One set was analyzed for sequence information by Sanger sequencing and one set was expanded for cell line generation. Four clones were chosen for experiments reported in this publication. Two wild-type (WT) clones, and two *SORL1* KO clones were selected. Sequencing data for all cell lines was confirmed by measuring protein expression using western blot techniques as described in our previous studies (Knupp et al., 2020; Mishra et al., 2023; Mishra et al., 2022). All four clones were shown to have normal karyotypes. Authentication by Sequencing data confirming CV cell lines by presence of a SNP unique to this genetic background is also described in our previous study(Knupp et al., 2020; Mishra et al., 2023; Mishra et al., 2022) . All cell lines are routinely karyotyped by Diagnostic Cytogenetics, Inc. (Seattle, WA), and tested for mycoplasma (MycoAlert). CRISPR/Cas9 gRNA, ssODN, and Primer Sequences gRNA:ATTGAACGACATGAACCCTC ssODN: GGGAATTGATCCCTATGACAAACCAAATACCATCTACATTGAACGACATGAACCCTC TGGCTACTCCACGTCTTCCGA AGTACAGATTTCTTCCAGTCCCGGGAAAACCAGGAAG Forward primer: ctctatcctgagtcaaggagtaac Reverse primer: ccttccaattcctgtgtatgc PCR amplifies 458 bp sequence.

### 2.2 Differentiation of human iPSCs into microglia like cells (hMGLs)

hiPSCs were differentiated into hMGLs as previously described with some modifications (McQuade et al., 2018). Briefly, hiPSCs were plated in mTESR plus medium supplemented with ROCK Inhibitor (Y-27632; # A3008; Apex Bio) on Matrigel (Growth factor reduced basement membrane matrix; # 356231; Corning) coated 6 well plates (#657160; CELLSTAR). To begin hematopoietic progenitor differentiation, hiPSC aggregates were plated at a density of 20-40 aggregates per well of a 6 well plate which resulted in attachment of ∼10 colonies per well of a 6 well plate for all the cell lines. In our hands, this plating density protocol resulted in maximum yield and efficient differentiation of HPCs from hiPSCs without generation of cell debris. On day 0, mTESR plus medium was replaced with STEMdiff™ Hematopoietic Supplement A medium from the STEMdiff™ Hematopoietic kit (# 05310; STEMCELL technologies). On day 3, when colonies became flattened, medium was replaced with STEMdiff™ Hematopoietic Supplement B medium from the STEMdiff™ Hematopoietic kit (# 05310; STEMCELL technologies). Cells remained in this medium for 7 additional days. By day 12, non-adherent hematopoietic progenitor cells (HPCs) coming off from the flattened colonies were harvested by removing medium. Any remaining HPCs/floating cells were collected by gentle PBS washes. At this point, HPCs were either frozen using Bambanker cell freezing medium (#BBH01; Bulldog-Bio) or plated at a density of 0.2 million cells per well of a Matrigel coated 6 well plate in microglia differentiation medium for 24 days. Microglia differentiation medium comprised of DMEM-F12 (#11039047; Thermo Fisher Scientific), Insulin-transferrin-selenite (#41400045; Thermo Fisher Scientific), B27 (# 17504-044; Thermo Fisher Scientific), N2 (# 17502-048; Thermo Fisher Scientific), Glutamax (# 35050061; Thermo Fisher Scientific), non-essential amino acids (# 11140050; Thermo Fisher Scientific), monothioglycerol (# M1753; Sigma), Insulin (# I2643; Sigma) freshly supplemented with TGF-β (#130-108-969, Miltenyi), IL-34 (# 200-34; Peprotech) and M-CSF (#PHC9501; Thermo Fisher Scientific). On day 24, this medium was supplemented with CD200 (#C311; Novoprotein) and CX3CL1 (#300-31; Peprotech) for maturation of microglia. Cells remained in this medium for 8 days. On day 32, microglia differentiation was complete, and these cells were plated on Poly-l-lysine (#P6282; Sigma; 100µg/ml) coated coverslips (12mm diameter, #1760- 012; cglifesciences) in a 24 well plate for immunocytochemistry with appropriate antibodies or Poly-l-lysine coated 48 well plates for flow cytometry.

### 2.3 Meso-Scale Discovery (MSD) V-PLEX platform to measure extracellular levels of cytokines in hMGLs

Pro-inflammatory cytokines in the media were measured using the Proinflammatory Panel 1 (human) kit (#K15049D; MSD V-PLEX MULTI-SPOT assay System; MesoScale Diagnostics). For this assay, hMGLs were plated at a density of 100,000 cells/well of a matrigel coated 96 well plate and allowed to settle for 24 hours. Cells were then treated with 20 ng/ml IFNγ or 100ng/ml Lipopolysaccharide (LPS) for 24 hours. Conditioned media was harvested at the end of 24 hours and stored at -80°C until use. 50µl of media was used to run the MSD V-plex assay as per manufacturer’s protocol and MSD Quick plex SQ120 instrument was used to detect analytes. This MSD V-Plex Proinflammatory Panel detects a panel of Pro-inflammatory cytokines including IL- 1β, IL-6, IL-8, IFN-γ, TNF-α, IL-13, IL-2, IL-4, IL-10 and IL-12p70, known to be altered in AD patients (Park et al., 2020; Taipa et al., 2019).

### 2.3 ELISAs to measure extracellular levels of cytokines in hMGLs

Extracellular levels of the cytokines, IL-6 and IL-1β were measured using the human IL-6 ELISA kit (#ab178013, abcam) and human IL-1β ELISA kit (#BMS224-2; Thermo Fisher Scientific). To test secretion of these cytokines upon activation with the lysosomal exocytosis activator, calcimycin, cells were treated with 1µM calcimycin for 24hours, extracellular levels (media) and intracellular levels (lysates) of IL-1β and IL-6 were measured using the ELISA kits described above.

### 2.4 In vitro fibrillation of Amyloid-β (1-42) (fAβ)

HiLyte Fluor 647 beta-amyloid (1-42) (# AS-64161, Anaspec) was fibrillized as described previously (Amos et al., 2017). Briefly, Aβ-42 was resuspended in sterile 1X PBS to generate a final concentration of 100µM, placed at 37°C for 5 days for fibrillization, aliquoted and stored at -80°C.

### 2.5 Phagocytosis of fibrillar Amyloid-β (1-42) and oligomeric Amyloid-β (1-42)

To measure phagocytosis of fAβ by hMGLs, Aβ labeled with a fluorophore, HiLyte Fluor 647 beta-amyloid (1-42) (# AS-64161, Anaspec) was fibrillized as described above and commercially available Oligomeric Aβ (1-42) (# SPR-488; Stress Marq Biosciences) was labeled with the Alexa Fluor™ 647 fluorophore using the Microscale Protein Labeling Kit (#A30009; Thermo Fisher Scientific)fAβ-647 (fAβ) and Oligomeric Aβ-647 (oAβ) were used for phagocytosis assay. hMGLs were plated at a density of 100,000 cells per well of a Poly-l-lysine (100µg/ml) coated 48 well plate. Cells were treated with this fAβ (0.1μM) and oligomeric Aβ (10µg/ml)for 30 mins, 5 hours and 24 hours. At the end of each time point, cells were washed with 1X PBS and harvested using accutase. Intracellular fluorescence of fAβ and oAβ was measured using flow cytometry. Data was analyzed using Flow Jo software. Fluorescence measured at 30 mins indicates uptake efficiency and the longer time points can either indicate sustained phagocytic capacity of hMGLs or accumulation of undegraded substrates in the lysosomes. As an additional method to test phagocytosis of fibrillar Aβ by hMGLs, phagocytosis assay was performed as described above. Subsequently, phagocytosis was examined using immunocytochemistry with cells fixed with 4 % PFA in PBS for 10 mins and imaged using the Leica SP8 microscope.

### 2.6 Phagocytosis of synaptosomes

To measure phagocytosis of synaptosomes, a phagocytosis assay was performed as previously described with some modifications(Beeken et al., 2022). Briefly, synaptosomes were harvested from Wildtype hiPSC-derived neurons using Syn-PER Synaptic Protein Extraction Reagent (# 87793; Thermo Fisher Scientific). 3 ml Syn-PER reagent was added to 1 10cm cell culture plate of neurons (∼30M cells), cells were harvested by scraping and centrifuged at 1200g for 10 mins. The supernatant was collected and centrifuged at 15,000g for 20 mins at 4°C and the pellet which are the synaptosomes (crude synaptosome extraction) were either resuspended in Krebs ringer solution (with 1X protease inhibitor) and frozen at -80°C or labeled with CM-Dil cell tracker dye (final concentration 3μM) as per manufacturer’s instructions (#C7000; Thermo Fisher Scientific). CM-Dil labeled synaptosomes were added to hMGLs plated on Poly-l-lysine coated 48 well plates and cells were harvested at the end of different time points including 30 mins, 5 hours, and 24 hours. The intracellular fluorescence intensity of CM-Dil labeled synaptosomes was measured using flow cytometry as a readout of phagocytosis of synaptosomes by hMGLs. To further confirm phagocytosis of synaptosomes by hMGLs, phagocytosis assay was performed as described above followed by immunocytochemistry with cells fixed with 4 % PFA in PBS for 10 mins and imaged using the Leica SP8 microscope.

### 2.7 Uptake assays

To test the uptake of FITC-Dextran (0.5mg/ml#46944-100MG-F; Millipore Sigma) and Transferrin from Human Serum, Alexa Fluor™ 647 Conjugate (25µg/ml; #T23366; Thermo Fisher Scientific), hMGLs were treated with each substrate for 2 hours in a cell culture incubator and fluorescence intensity was measured using flow cytometry. Analysis was performed using Flow Jo software.

### 2.8 Western blotting

For western blot analysis, cell lysates were harvested from hMGLs using RIPA protein lysis buffer (#20-188; Millipore) containing 1X protease and 1X phosphatase inhibitors (#PI78443; ThermoFisherScientific). Total protein concentration was quantified using Pierce BCA assay kit (#23225; Thermo Fisher Scientific). 10-20μg of protein lysates were run on 4%–20% Mini-PROTEAN TGX Precast Protein Gels (#4561096; Biorad) and transferred to PVDF membranes. Membranes were blocked with 5% w/v non-fat dry milk in 1X PBS (blocking buffer), washed with 1X PBS with 0.05% tween 20 (PBST) and incubated overnight at 4°C with the following primary antibodies diluted in 0.2% sodium azide+5% BSA in 1X PBS : Sortilin-related receptor 1 (SORLA) at 1:1000 (# ab190684; abcam); Mouse monoclonal anti-Actin (clone A4) (#MAB1501; Millipore Sigma) at 1:2000; Rabbit monoclonal anti-HEXB (#ab140649; abcam); Goat anti-Cathepsin B (#ab214428; abcam); Mouse monoclonal anti-Cathepsin D (#ab6313; abcam) and Rabbit monoclonal anti-LAMP1 (#9091S; Cell signaling); Rabbit Polyclonal P2Y6 receptor (#APR-106; Alomone labs); Human TREM2 (#MAB17291; R&D) and Anti-M6PR (cation independent) antibody [EPR6599] (#ab124747; Abcam). Membranes were then washed with PBST, incubated in corresponding HRP conjugated secondary antibodies for 1h. The following secondary antibodies were used. Goat anti-Mouse IgG (H+L) Secondary Antibody HRP (#31430; Thermofisher Scientific) Goat anti-Rabbit IgG (H+L) Secondary Antibody HRP (# 31460; Thermofisher Scientific) and Donkey anti-goat IgG (H+L) Secondary Antibody (#A15999; ThermoFisher Scientific). Membranes were then washed thrice with PBST, probed with either Clarity Western ECL substrate (#1705061; Biorad) or Clarity Max Western ECL Substrate and scanned using an Odyssey Clx imaging system (Li-Cor) to detect bound proteins on membranes. To normalize protein levels in the media, ponceau staining was used.

### 2.9 Lysosomal degradation assay

Lysosomal degradation in hMGLs was measured using DQ-BSA (#12051; Thermo Fisher Scientific) which is a fluorogenic substrate for proteases and fluoresces only when degraded in lysosomes. For this assay, hMGLs were plated at a density of 100,000 cells per well of a Poly-l-lysine coated 48 well plate. A pulse-chase experiment was performed wherein hMGLs were treated with 20µg/ml DQ-BSA for 20 mins. At the end of this timepoint, cells were washed with 1X PBS and cultured in complete microglia medium for 30 mins, 2 hours, and 5 hours. At the end of each timepoint, cells were washed with 1X PBS and harvested using accutase. Intracellular fluorescence intensity of DQ-BSA was measured using flow cytometry and data was analyzed using flow jo software. Altered fluorescent intensity of DQ-BSA was measured as a readout of altered lysosomal degradation. As an additional method to examine lysosomal degradation, we performed pulse chase experiment as described above and immunocytochemistry (cells fixed after 2h) after treating cells with DQ-BSA and intracellular fluorescence intensity was noted as a readout of lysosomal degradation. Treatment with the lysosomotropic agent chloroquine (50μM) was used as a negative control for both experiments.

### 2.10 Lysosomal enzyme activity assays

To measure lysosomal enzyme function, lysosomal enzyme activity assays were performed. For hMGLs generated using protocol based on McQuade et al 2018, 200,000 hMGLs were plated per well of a poly-l-lysine coated 48 well plate. Cells were allowed to settle for 24 hours. Specifically, enzyme activity of lysosomal enzymes, Cathepsin D (#ab65302; abcam), Cathepsin B (#ab 65300; abcam) and Hexosaminidase B (#MET-5095; Cell Bio labs) was measured as per manufacturers protocol.

### 2.11 Immunocytochemistry

hMGLs were seeded at a density of 200,000 cells per well of a Poly-l-lysine coated 24-well plate on glass coverslips. After 2 days in culture, cells were fixed in 4% paraformaldehyde (PFA, Alfa Aesar, Reston, VA) for 10 minutes. Cells were washed with 1X PBS + 0.05% tween 20 detergent, incubated in blocking buffer containing 5% goat (#005-000-121; Jackson ImmunoResearch) or donkey serum (#017-000-121; Jackson ImmunoResearch), permeabilized using 0.1% Triton X- 100 (Sigma Aldrich, St Louis, MO) for 15 minutes at room temperature and then incubated in a primary antibody dilution in blocking buffer overnight at 4°C. Cells were then washed three times with PBS + 0.05% tween 20 and incubated with a secondary antibody dilution in blocking buffer for 1 hour at room temperature in the dark. Cells were then washed with 1X PBS with tween 20 and mounted on glass slides with ProLong Gold Antifade mountant with DAPI(#P36931; Thermo Fisher Scientific). To measure surface expression, cells were not permeabilized. Surface expression was reported as fluorescence intensity of surface proteins normalized to cell area as measured by the Image J software.

### 2.12 Colocalization analysis

To investigate colocalization of lysosomal enzymes with the TGN or lysosomes, hMGLs were plated in a 24 well plate with 12mm diameter coverslips, allowed to settle for 48 hours and fixed with 4% PFA in PBS. Immunocytochemistry was performed using primary antibodies against lysosomal enzymes, Cathepsin B, Cathepsin D, HEXB, Cation-Independent Mannose-6 phosphate receptor (CIMPR) and TGN marker, TGN38 and late endosomal and lysosomal marker, LAMP1. Cells were co-labelled for either TGN and lysosomal enzymes or LAMP1 and lysosomal enzymes. For the lysosomal enzymes and CIMPR, the same antibodies were used for both immunocytochemistry and western blotting. To label TGN and lysosomes, TGN38 (B6) mouse monoclonal antibody (#166594; SantaCruz) and LAMP1 mouse monoclonal antibody (#sc20011; Santacruz) were used. A minimum of 10 fields of confocal images were captured using the Leica SP8 microscope. Images were analyzed using Image J software. Median filtering was used to remove noise from images and Otsu thresholding was applied to all images. Colocalization was quantified using the JACOP plugin in ImageJ software and presented as Mander’s correlation coefficient.

### 2.13 Confocal microscopy and image processing

All microscopy and image processing were performed under blind conditions. Z stack images were acquired using the Leica TCS SP8 confocal laser microscope with 63x/1.40 oil lens (Leica Microsystems) and modified/processed using Leica Application Suite X 3.5.5.19976 with LIGHTNING adaptive deconvolution (Leica Microsystems). 10-20 fields were imaged per experiment per clone, which included a total of 50-100 cells. For some experiments, images were processed further using Image J software (Schindelin et al., 2012).

### 2.14 Measurement of surface LAMP1 expression

To measure surface LAMP1 expression, hMGLs were plated at a density of 100,000 cells per well of a 48 well plate and allowed to settle for 48 hours. Cells were then either treated with the calcium ionophore, calcimycin (#C4403; Sigma) +4mM CaCl2+0.1% BSA in PBS to stimulate LE or 0.1%BSA in PBS as control and placed in the cell culture incubator for 30 mins. Next, cells were placed on ice for 15 mins and treated with cold Mouse monoclonal antibody against LAMP1(#328611; Biolegend) specific to the N terminal luminal epitope of the antigen, for an hour on ice. Cells were then washed with 1X PBS and either treated with accutase and used for flow cytometry or fixed with 4% PFA in PBS for 5 mins and used for immunocytochemistry. Surface expression of LAMP1 was presented as fluorescence measured by flow cytometry and Flow Jo software or immunocytochemistry and Image J software.

### 2.15 Quantification and statistical analysis

All data presented in this study represent multiple hiPSC clones. This includes two clones for WT hMGLs and two clones for *SORL1* KO hMGLs. All data represent at least six independent experiments (called replicates) per clone. Experimental data were tested for normal distributions using the Shapiro-Wilk normality test. Normally distributed data were analyzed using parametric two-tailed unpaired t tests, one-way ANOVA tests, or two-way ANOVA tests. Significance was defined as a value of p < 0.05. All statistical analysis was completed using GraphPad Prism software.

## 3 Results

### 3.1 Loss of *SORL1* decreases lysosomal degradation in hMGLs

We have previously shown that *SORL1* KO hMGLs have enlarged lysosomes indicating lysosome dysfunction and stress (Mishra, 2024). We hypothesized that lysosomal dysfunction would impair the degradative capacity of this organelle. To test this hypothesis, we generated WT and *SORL1* KO hematopoietic progenitor cells (HPCs) and hMGLs using a previously published protocol (McQuade et al., 2018) with minor modifications (described in the methods section). HPC-specific expression was confirmed in WT and *SORL1* KO HPCs using immunostaining with the HPC marker CD43 and microglia specific expression was demonstrated by immunostaining with the microglia markers IBA1, and CX3CR1. Loss of *SORL1* did not alter differentiation of hiPSCs into HPCs and hMGLs (Figure S1). To determine if overall proteolytic lysosomal degradation is affected, we performed a pulse chase assay with DQ-Red-BSA, a fluorogenic substrate that fluoresces only upon degradation (Marwaha & Sharma, 2017) (Figure 1A). Mean fluorescence intensity (MFI) was measured by flow cytometry as a readout of lysosomal degradation. Both WT and *SORL1* KO hMGLs showed an increase in MFI over a period of 2h suggesting our hiPSC-derived microglia were capable of lysosomal degradation *in vitro*, although *SORL1* KO hMGLs demonstrated reduced MFI of DQ-Red-BSA as compared to WT hMGLs at all time points tested (Figure 1B), suggesting either decreased uptake and/or degradation of the substrate. Inhibiting lysosome proteolysis with the lysosomotropic reagent chloroquine abrogated lysosomal degradation of DQ-Red-BSA in both WT and *SORL1* KO hMGLs (Figure 1B), validating our degradation assays. To test if decreased fluorescence intensity of DQ-BSA observed in *SORL1* KO hMGLs is caused by decreased uptake of the reagent, we tested uptake of a proxy reagent, BSA conjugated to a red fluorophore (BSA-594), which also undergoes pinocytic uptake similar to DQ-BSA. A proxy reagent was used because short-term uptake of DQ-BSA cannot be measured directly since it fluoresces only upon reaching the lysosomes. Interestingly, we observed an increased uptake of BSA-594 in *SORL1* KO hMGLs as compared to WT hMGLs (Figure 1C). To further corroborate our data, we performed an uptake assay with a general pinocytosis marker, FITC-Dextran and observed a similar trend in *SORL1* KO hMGLs (Figure S4A). Together, this data suggests that *SORL1* KO hMGLs have reduced lysosomal degradation of DQ-Red-BSA rather than reduced uptake of substrates. To further confirm our findings with an independent method, we performed a pulse chase assay and immunocytochemistry. We incubated hMGLs with DQ-Red-BSA similar to the experiment described above and cells were fixed after 2 hours. Immunocytochemistry was performed to label cells with the microglia marker CX3CR1. In support of our hypothesis, we observed decreased lysosomal degradation in *SORL1* KO hMGLs as compared to WT hMGLs (Figure 1D-F). Overall, our data demonstrated that loss of *SORL1* causes reduced lysosomal degradation in hMGLs.

**Figure 1.**
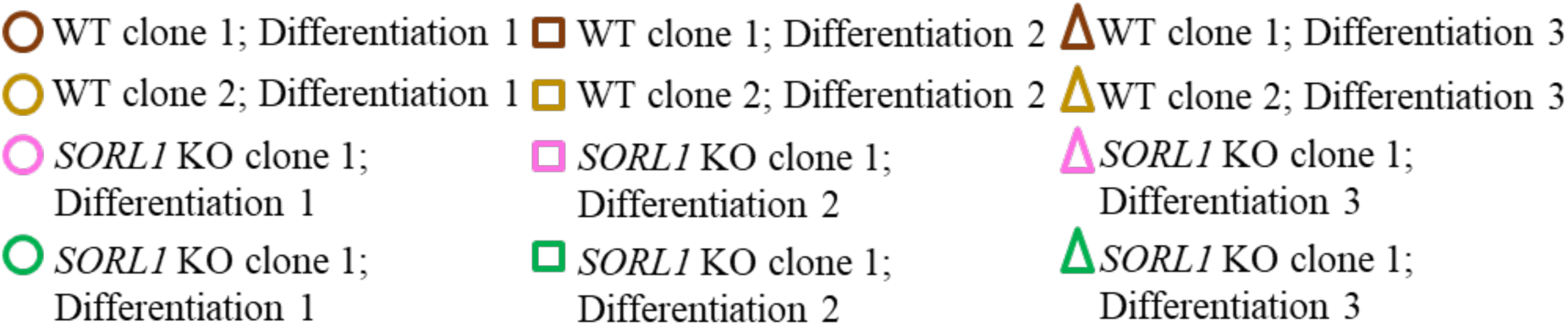
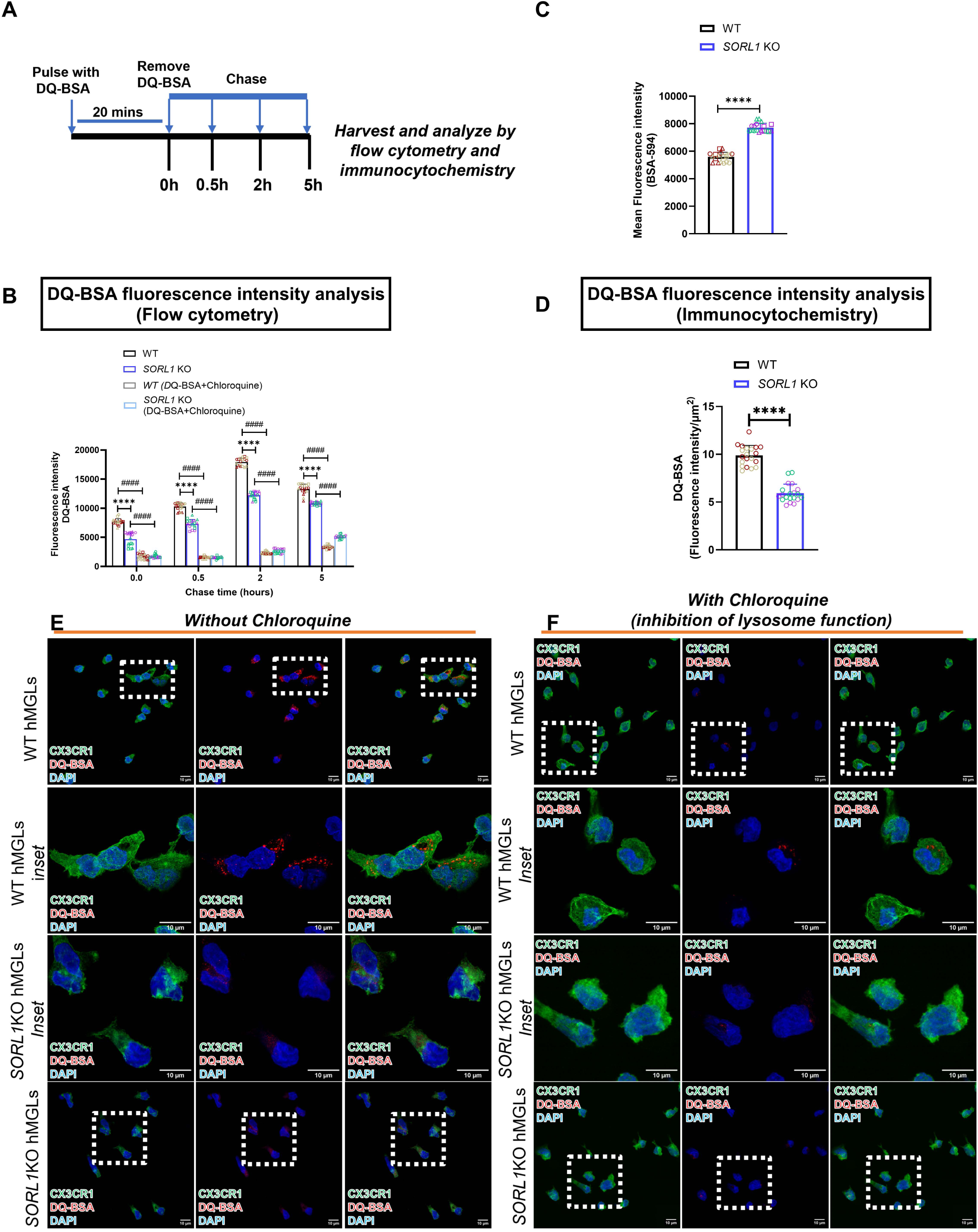
Loss of *SORL1* results in decreased lysosomal degradation in hMGLs. (A) Schematic illustrating experimental design using DQ-BSA, a fluorogenic protease substrate that fluoresces only upon lysosomal degradation by proteases. (B) A pulse chase experiment performed with WT and *SORL1* KO hMGLs and Mean fluorescence intensity (MFI) measured using flow cytometry. *SORL1* KO hMGLs show decreased MFI of DQ-BSA at all timepoints indicating decreased lysosomal degradation. The lysosomotropic agent chloroquine that inhibits lysosomal degradation was used as control. Chloroquine treatment shows decreased lysosomal degradation of DQ-BSA in both WT and *SORL1* KO hMGLs. (C) Flow cytometry showing increased MFI of BSA-594 in *SORL1* KO hMGLs relative to WT hMGLs suggestive of increased uptake of BSA in hMGLs with loss of *SORL1*. (D-F) Immunocytochemistry of hMGLs treated with DQ-BSA using the experimental design in (A), cells fixed for 2h and labeled with the microglia marker, CX3CR1 showing decreased lysosomal degradation in *SORL1* KO hMGLs as compared to WT hMGLs, validating results obtained from the experiments using flow cytometry. Similar to the flow cytometry experiment, chloroquine was used as a control and showed decreased MFI in both WT and *SORL1* KO hMGLs indicating reduced lysosomal degradation Quantification of fluorescence intensity of DQ-BSA using Image J software after immunocytochemistry with microglia marker, CX3CR1 and treatment with DQ-BSA. (E-F) I. 2 isogenic clones per genotype (WT and *SORL1* KO) and 9 independent replicates (3 differentiations and 3 technical replicates per differentiation) per clone per genotype (N=18 independent replicates) were used for these experiments. Data represented as mean ± SD and analyzed using parametric two-tailed unpaired t test and 2-way ANOVA with Tukeys multiple comparison test. Significance while comparing WT to *SORL1* KO hMGLs was defined and depicted as a value of *p < 0.05, **p < 0.01, ***p < 0.001, and ****p < 0.0001, ns=not significant and while comparing different treatments of each genotype was defined and depicted as a value of #p < 0.05, ##p < 0.01, ###p < 0.001, and ###p < 0.0001, ns=not significant.

### 3.2 Loss of *SORL1* decreases intracellular lysosomal enzyme activity in hMGLs

Next, we investigated intracellular lysosomal enzyme activity as a potential cause for altered lysosomal degradation observed in *SORL1* KO hMGLs. We tested intracellular enzyme activity of three key lysosomal enzymes, Beta-hexosaminidase subunit beta (HEXB), Cathepsin D (CTSD) and Cathepsin B (CTSB). All three enzymes are highly expressed in microglia, CTSD and CTSB have been proposed to degrade Aβ and tau, and altered expression of all three enzymes has been observed in AD(Cermak et al., 2016; Di Spiezio et al., 2021; Hamazaki, 1996; Ii et al., 1993; Kenessey et al., 1997; Kuil et al., 2019; Sierksma et al., 2020; Sun et al., 2008; Whyte et al., 2022). Furthermore, altered expression or activity of these enzymes causes significant lysosome dysfunction and affects Aβ deposition in mouse brains (Keilani et al., 2012; Koike et al., 2000; Oberstein et al., 2021; Suire et al., 2020; Suzuki et al., 2022; Wang et al., 2012). Depletion of *SORL1* resulted in reduced lysosomal enzyme activity of HEXB, CTSD and CTSB (Figure 2A-C) without altering protein expression in *SORL1* KO hMGLs relative to WT hMGLs (Figure 2 D-I). Chloroquine treatment was used as a negative control in enzyme activity assays and, as expected, resulted in decreased lysosomal enzyme activity for all the enzymes (Figure 2A-C). To further confirm our hypothesis and account for potential heterogeneity introduced by microglia differentiation protocols, we tested lysosomal enzyme activity in hiPSC-derived microglia generated using a different protocol (Dolan et al., 2023). Characterization of these hMGLs showed no difference in expression of microglia-specific markers between WT and *SORL1* KO cells (Figure S2A). Using these hMGLs, we independently confirmed reduction in intracellular CTSD activity (Figure S2B), validating our hypothesis that loss of *SORL1* results in reduced lysosomal enzyme activity. Overall, our data suggests that decreased lysosomal enzyme activity, rather than changes in overall lysosomal protein expression, contributes to the reduction in lysosomal degradation observed in *SORL1* KO hMGLs. Our results are concordant with a recent study that showed no difference in mRNA and protein expression of CTSD, CTSB and HEXB in SORL1 KO hMGLs using mRNA-sequencing and TMT proteomics (Lee et al., 2023). Furthermore, lysosomal phenotypes due to *SORL1* deficiency are not due to differences between *in vitro* differentiation protocols.

**Figure 2.**
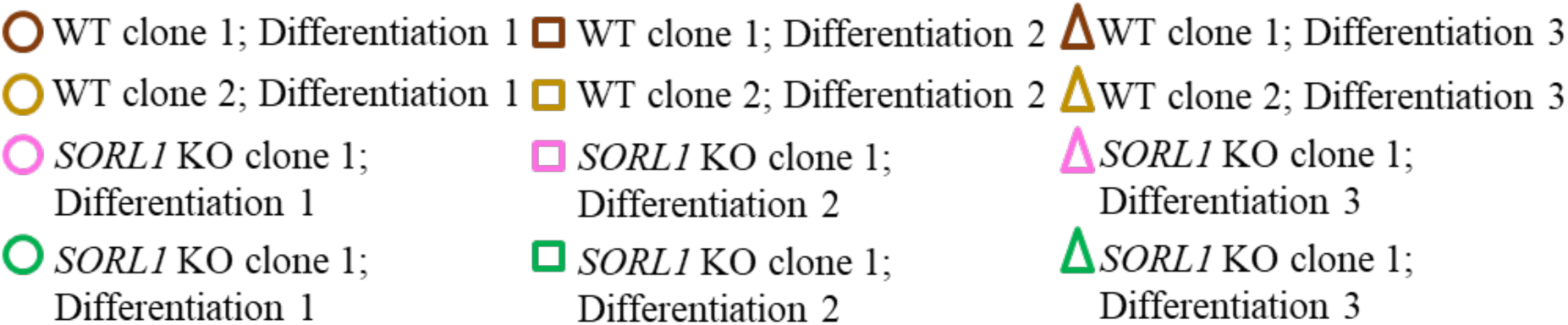
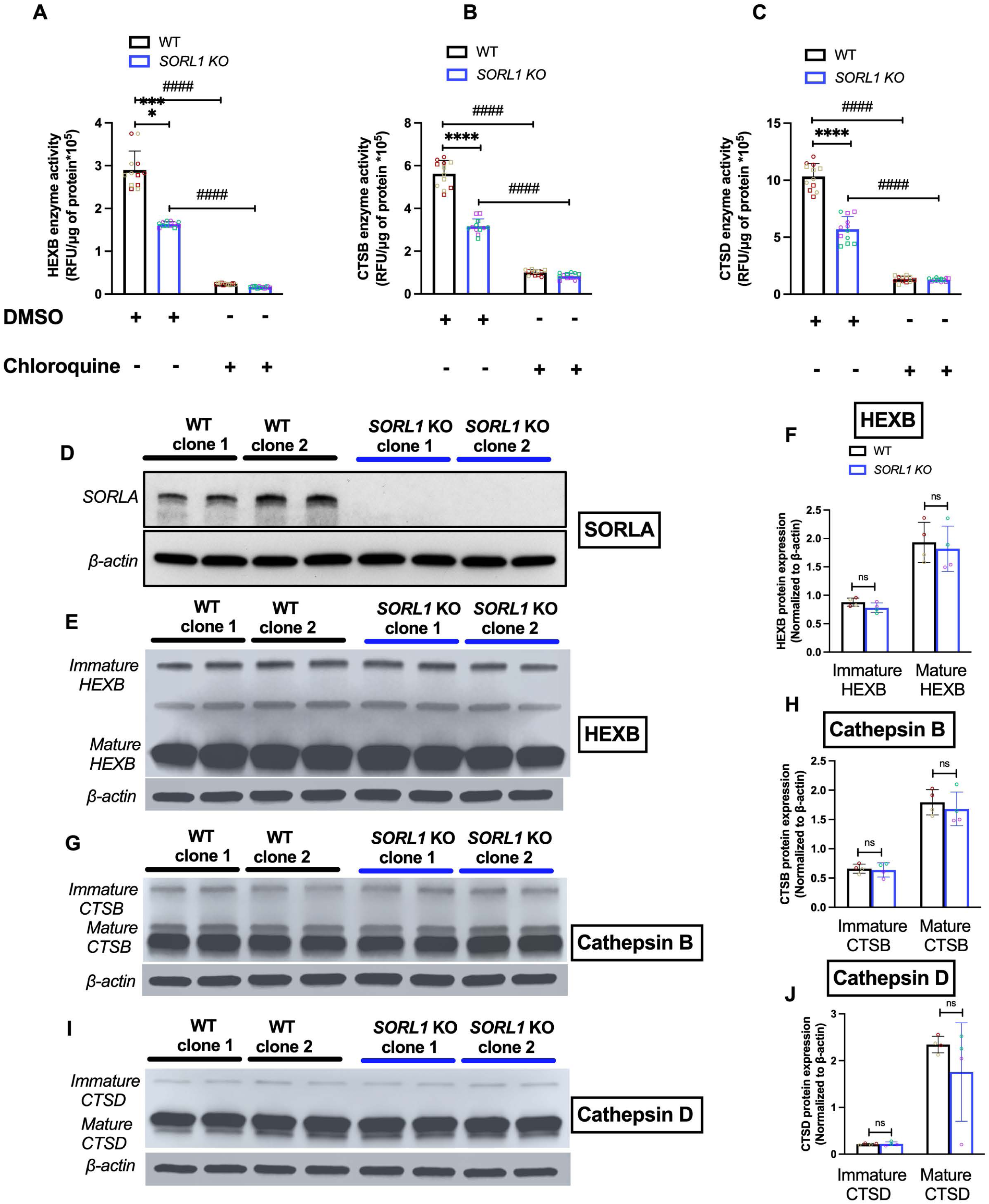
Loss of *SORL1* results in decreased lysosomal enzyme activity in hMGLs. (A-C) Enzyme activity assays based on fluorometric detection show reduced enzyme activity of lysosomal enzymes (A) HEXB, (B) Cathepsin B and (C) Cathepsin D in *SORL1* KO hMGLs as compared to WT hMGLs. Both WT and *SORL1* KO hMGLs show reduced enzyme activity of all the enzymes tested when treated with the lysosome function inhibitor, chloroquine, which was used as a control. (D-I) Western blotting demonstrates loss of SORLA protein expression (D) but no change in total protein levels of immature and mature forms of HEXB (E-F), Cathepsin B (G- H) and Cathepsin D (I-J) in *SORL1* KO hMGLs relative to WT hMGLs. For the enzyme activity assays, 2 isogenic clones per genotype (WT and *SORL1* KO) and 6 independent replicates (2 differentiations and 3 technical replicates per differentiation) per clone per genotype (N=12 independent replicates) were used and for the western blot experiments, 2 isogenic clones per genotype (WT and *SORL1* KO) and 2 independent replicates (1 differentiation and 2 technical replicates) per clone per genotype (N=4 independent replicates) were utilized. Data represented as mean ± SD and analyzed using parametric two-tailed unpaired t test and 2-way ANOVA with Tukeys multiple comparison test. Significance while comparing WT to *SORL1* KO hMGLs was defined and depicted as a value of *p < 0.05, **p < 0.01, ***p < 0.001, and ****p < 0.0001, ns=not significant and while comparing different treatments of each genotype was defined and depicted as a value of #p < 0.05, ##p < 0.01, ###p < 0.001, and ###p < 0.0001, ns=not significant

### 3.3 Loss of *SORL1* results in altered trafficking of lysosomal enzymes in hMGLs

Newly synthesized lysosomal enzymes are transported from the trans Golgi network (TGN) to the lysosome using either a mannose-6-phosphate (M6P)-dependent pathway primarily regulated by Cation Independent Mannose-6-phosphate receptor (CIMPR) and the multi-protein complex retromer (Arighi et al., 2004; Thomas Braulke & Juan S. Bonifacino, 2009; Cui et al., 2019) or in a M6P-independent pathway in which enzymes are directly targeted to the lysosome(Gauthier et al., 2024; Hasanagic et al., 2015). SORLA can bind to the retromer complex (Fjorback et al., 2012) for various sorting functions and the retromer complex associates with CIMPR to regulate sorting of lysosomal enzymes (Cui et al., 2019; Nielsen et al., 2007). Thus, we hypothesized that loss of SORLA would result in altered trafficking of lysosomal enzymes from the TGN to the lysosomes. To test whether lysosomal enzymes are intracellularly mis-localized with loss of *SORL1*, we used immunocytochemistry with antibodies that recognized both immature and mature forms of the lysosomal enzymes Cathepsin B, Cathepsin D and HEXB. We measured colocalization of these enzymes at the TGN and late endosomes and lysosomes using the TGN marker TGN38 and the late endosome and lysosome marker, LAMP1. Our results showed increased colocalization of all three enzymes with TGN38 (Figure 3A,C,E) and decreased colocalization with LAMP1 (Figure 3B,D,F) suggesting that the enzymes are not appropriately trafficked to lysosomes from the TGN. As we observed decreased enzyme activity but no change in total protein expression of CTSD, CTSB and HEXB, our data suggested that dysfunction in lysosomal degradation is likely due to mis-trafficking of lysosomal enzymes.

**Figure 3.**
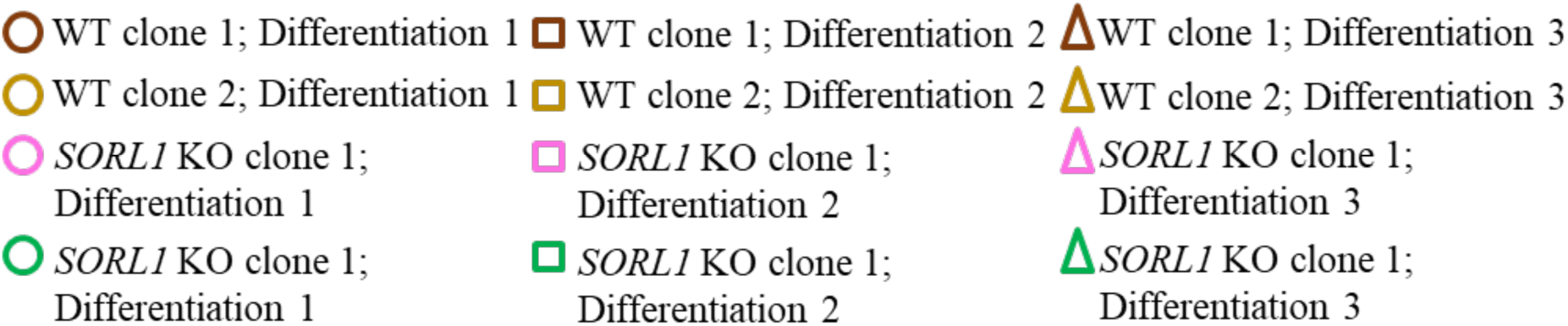
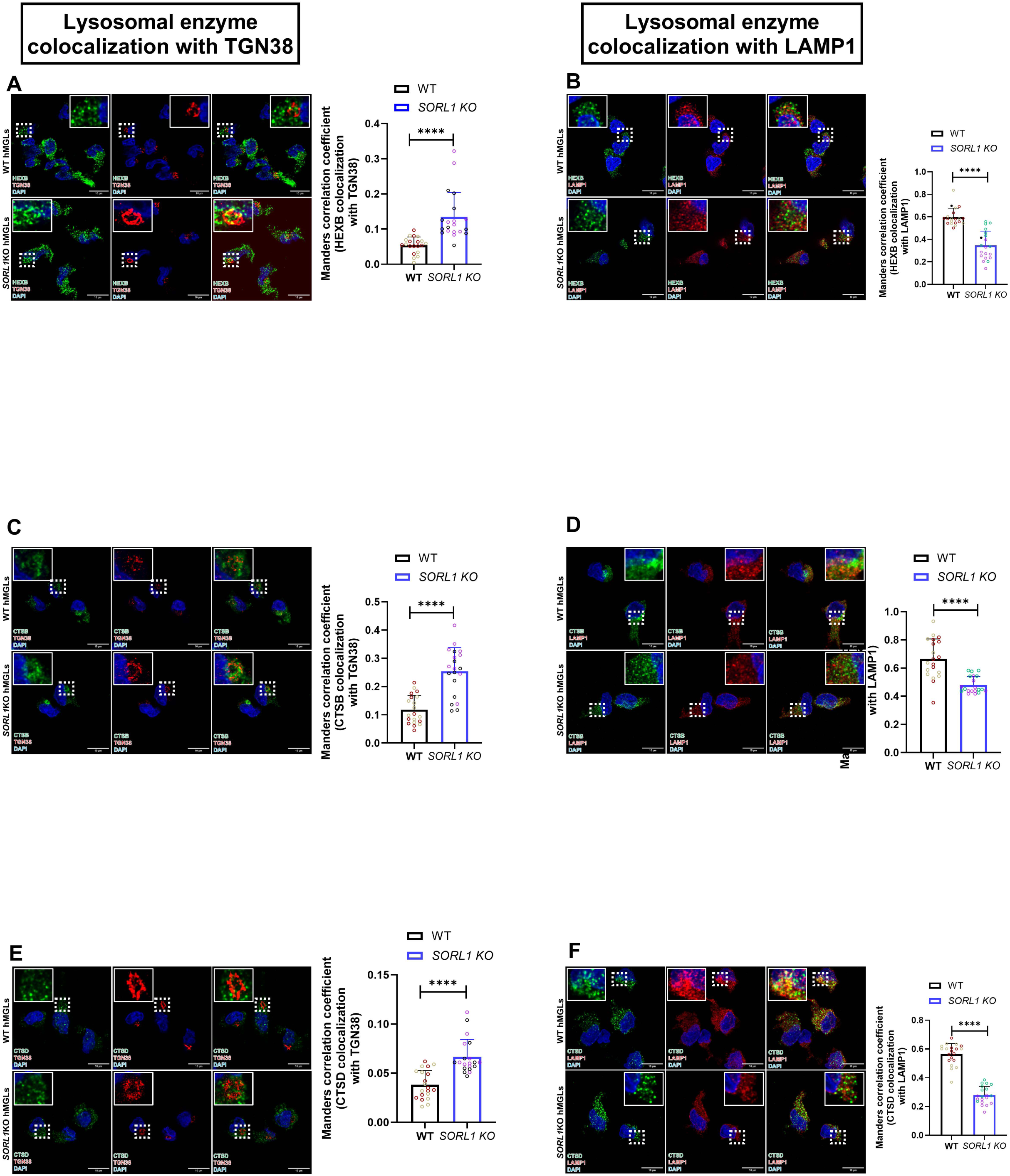
Loss of *SORL1* results in altered trafficking of lysosomal enzymes in hMGLs. Immunocytochemistry and colocalization analysis used to measure colocalization of lysosomal enzymes HEXB, Cathepsin B and Cathepsin D with TGN and lysosomes using TGN38 and LAMP1 antibodies respectively. As compared to WT hMGLs, *SORL1* KO hMGLs show increased colocalization of all the enzymes with TGN38 (Figure 3A,C,E) and decreased colocalization of all enzymes with LAMP1 (Figure 3B,D,F) suggestive of altered trafficking of lysosomal enzymes from TGN to lysosomes. Colocalization was measured using JACOP plugin in Image J software and presented as Manders correlation coefficient. Scale bar=10μm. 2 isogenic clones per genotype (WT and *SORL1* KO) and 10 images per clone per genotype (N=20 independent replicates) were used for these experiments. Each image comprised of at least 5-10 cells and hence a total of 50- 100 cells per clone per genotype were analyzed for colocalization analysis. Data represented as mean ± SD and analyzed using parametric two-tailed unpaired t test. Significance while comparing WT to *SORL1* KO hMGLs was defined and depicted as a value of *p < 0.05, **p < 0.01, ***p < 0.001, and ****p < 0.0001, ns=not significant.

### 3.4 Loss of *SORL1* results in reduced protein levels and dysfunctional trafficking of CIMPR from the TGN to the lysosomes in hMGLs

To further investigate the mechanism that causes dysfunctional trafficking of lysosomal enzymes in *SORL1* KO hMGLs, we tested if loss of SORLA causes altered expression and/or intracellular localization of the lysosomal enzyme trafficking protein, CIMPR. We measured colocalization of CIMPR at the TGN, using the TGN marker TGN38, and at late endosomes and lysosomes using LAMP1. Our data showed that *SORL1* KO hMGLs had reduced colocalization of CIMPR with LAMP1 and no change in colocalization with TGN38 (Figure 4A-B), suggesting that loss of *SORL1* leads to impaired CIMPR-dependent trafficking pathway of lysosomal enzymes out of the TGN to lysosomes. Additionally, we observed reduced protein levels of CIMPR in *SORL1* KO hMGLs as compared to WT hMGLs by Western blot analysis (Figure 4 C-D). Overall, these data demonstrate that depletion of SORLA results in altered protein expression and intracellular localization of the trafficking protein CIMPR, contributing to dysfunctional trafficking of lysosomal enzymes observed in *SORL1* KO hMGLs.

**Figure 4.**
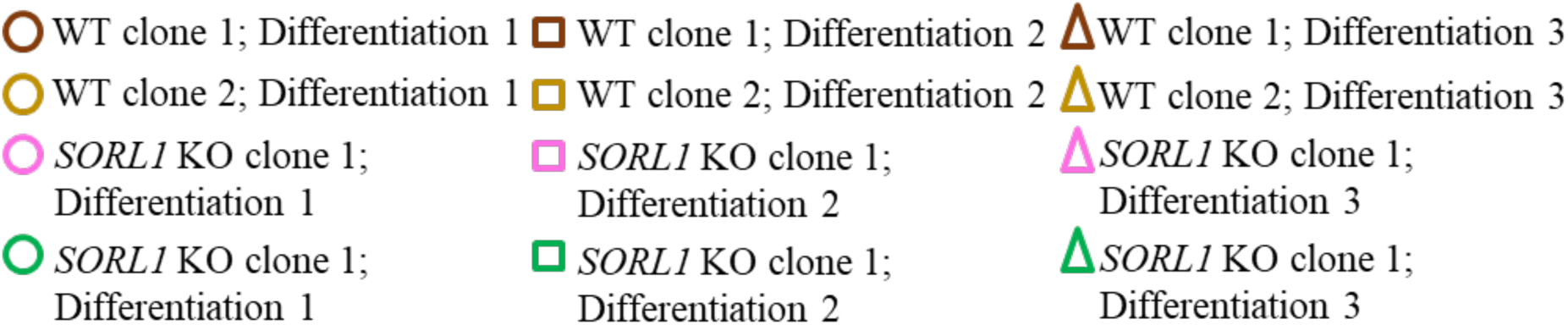
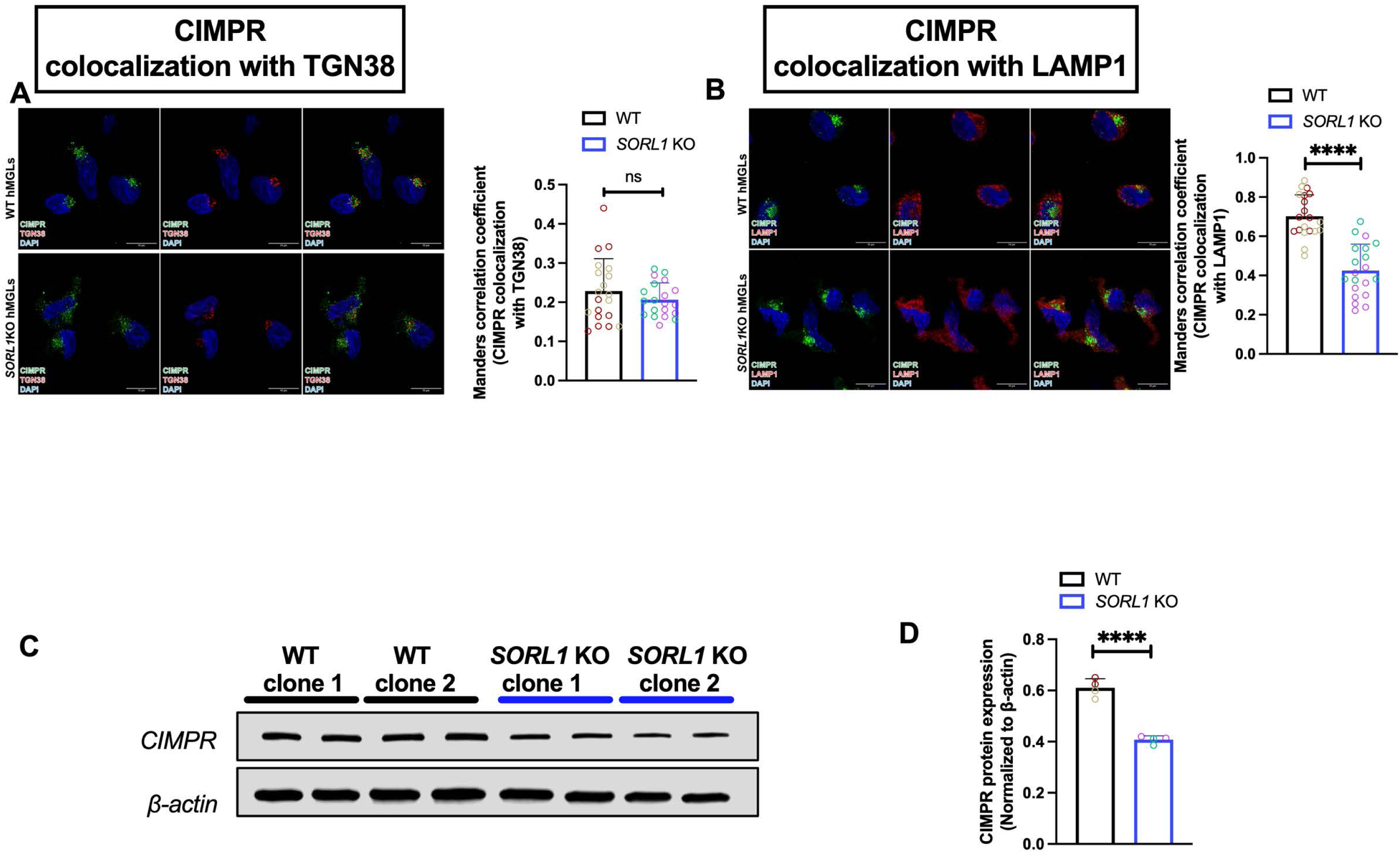
Loss of *SORL1* results in reduced protein levels and dysfunctional trafficking of Cation-independent Mannose-6-phosphate receptor (CIMPR), from the TGN to the lysosomes in hMGLs (A-B) Immunocytochemistry and colocalization analysis was used to measure colocalization of CIMPR with TGN and late endosomes and lysosomes using TGN38 and LAMP1 antibodies respectively. As compared to WT hMGLs, *SORL1* KO hMGLs show no change in colocalization of CIMPR with TGN38 (A) and decreased colocalization of CIMPR with LAMP1 (B). (C-D) Western blot and quantification showing decreased protein levels of CIMPR in SORL1 KO hMGLs as compared to WT hMGLs. Colocalization was measured using JACOP plugin in Image J software and presented as Manders correlation coefficient. Scale bar=10μm. 2 isogenic clones per genotype (WT and *SORL1* KO) and 10 images per clone per genotype (N=20 independent replicates; 1 differentiation) were used for these experiments. Each image comprised of at least 5-10 cells and hence a total of 50-100 cells per clone per genotype were analyzed for colocalization analysis. Data represented as mean ± SD and analyzed using parametric two-tailed unpaired t test. Significance while comparing WT to *SORL1* KO hMGLs was defined and depicted as a value of *p < 0.05, **p < 0.01, ***p < 0.001, and ****p < 0.0001, ns=not significant.

### 3.5 Loss of *SORL1* causes lysosomal accumulation of Aβ and synaptosomes in hMGLs

Given that depletion of *SORL1* causes reduced lysosomal degradation in hMGLs, we next hypothesized that phagocytosed substrates might accumulate in lysosomes in SORL1-deficient hMGLs. To test this idea, we measured lysosomal accumulation of fluorophore-labeled fibrillar Aβ 1-42 (fAβ) and CM-Dil dye labeled synaptosomes using pulse chase assays and immunocytochemistry. Consistent with our hypothesis, *SORL1* KO hMGLs showed increased accumulation of both substrates as evidenced by increased colocalization of Aβ 1-42 (Figure 5A-B) and synaptosomes (Figure 5C-D) with LAMP1. To further confirm that reduced lysosomal catabolism contributes to intracellular accumulation of both substrates, lysosome catabolism was inhibited by chloroquine and lysosomal accumulation of fAβ and synaptosomes was measured. As expected, chloroquine treatment caused a significant increase in lysosomal accumulation of both substrates in both WT and *SORL1* KO hMGLs (Figure 5B,D). Increased phagocytosis can contribute to increased accumulation of these substrates, so we tested whether phagocytic uptake was altered in *SORL1* KO hMGLs. To analyze uptake and phagocytosis we incubated cells with fluorophore-labeled fAβ and synaptosomes for 30 mins, 5h and 24h and observed increased uptake and sustained phagocytosis of both substrates in *SORL1* KO hMGLs as compared to WT hMGLs (Figure S4 D-G). Interestingly, we did not observe a difference in internalization of oligomeric Aβ (Figure S4C), consistent with previous reports(Ovesen et al., 2024). This suggests that *SORL1* may alter phagocytic uptake in a substrate-specific manner or that the dynamics of these process may be variable. Because we consistently observed increased uptake of fAβ and synaptosomes, we wondered whether loss of SORLA expression might alter cell surface localization of phagocytic receptors. To test this idea, we analyzed cell surface localization of two key phagocytic receptors known to be involved in phagocytosis of fibrillar Aβ and synaptosomes. TREM2 is known to phagocytose fibrillar Aβ and synaptosomes in microglia (La Rosa et al., 2023; McQuade et al., 2020) and the P2Y6 receptor is a potent phagocytic receptor involved in microglial phagocytosis of synaptosomes (Dundee & Brown, 2024; Dundee et al., 2023). We observed increased surface localization of both receptors in *SORL1* KO hMGLs (Figure S5 A-D) without alteration of total protein levels of TREM2 and P2Y6 receptor (Figure S5 E-F) suggesting that SORLA may play a role in maintaining the number of receptors at the cell surface required for phagocytosis and its absence causes aberrant localization of these receptors. We also tested uptake of a general endocytosis marker, transferrin conjugated to Alexa fluor 647 and also show increase uptake of this substrate in *SORL1* KO hMGLs (Figure S4 B). Overall, these data suggest that, in our hMGLs, SORLA may increase phagocytosis of fibrillar Aβ and synaptosomes by increasing surface localization of phagocytic receptors, TREM2 and P2Y6 in hMGLs. Together, our data demonstrate that through increased phagocytic uptake combined with impaired lysosomal degradation loss of *SORL1* significantly increases lysosomal dysfunction in human microglia.

**Figure 5.**
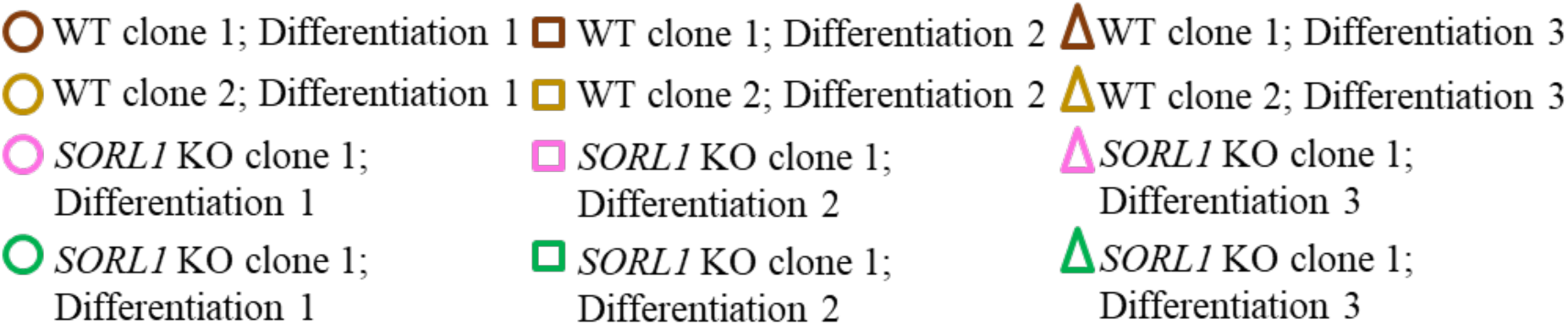
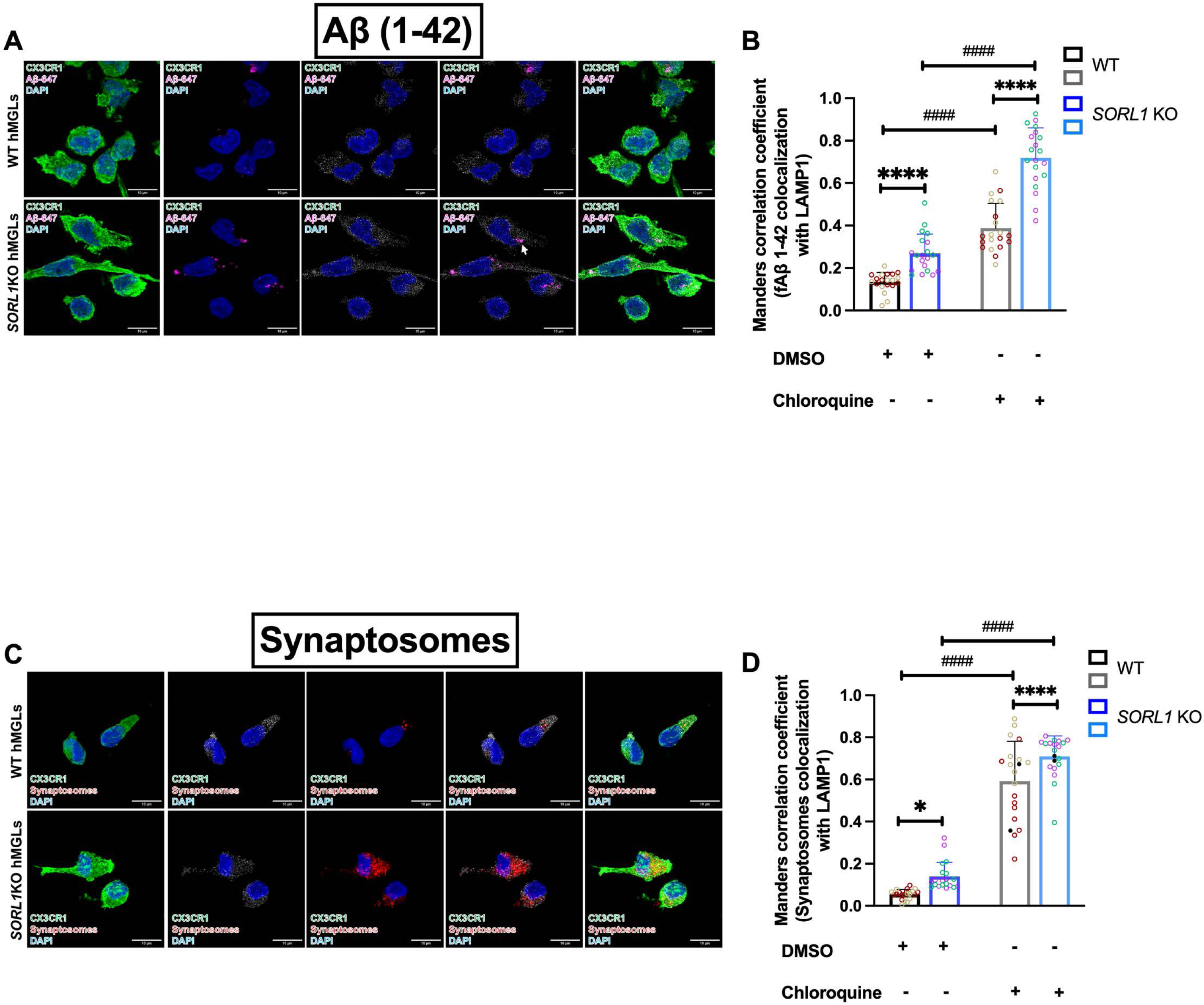
Loss of *SORL1* causes increased lysosomal accumulation of fibrillar Aβ 1-42 (fAβ) and synaptosomes in hMGLs. (A-D) Immunocytochemistry and colocalization analysis with and without treatment with chloroquine, showing increased colocalization of fAβ (A-B) and synaptosomes (C-D) with lysosome marker, LAMP1, in *SORL1* KO hMGLs as compared to WT hMGLs suggestive of increased lysosomal accumulation. Chloroquine treated WT and SORL1 KO hMGLs show increased lysosomal accumulation of both substrates as compared to their untreated counterparts. Colocalization analysis performed using JACOP plugin of Image J software and data is presented as Manders correlation coefficient. Scale bar=10μm. 2 isogenic clones per genotype (WT and *SORL1* KO) and 10 images per clone per genotype (N=20 independent replicates; 1 differentiation) were used for these experiments. Each image comprised of at least 5-10 cells and hence a total of 50-100 cells per clone per genotype were analyzed for colocalization analysis. Data represented as mean ± SD and analyzed using parametric two-tailed unpaired t test and 2-way ANOVA with Tukeys multiple comparison test. Significance while comparing WT to *SORL1* KO hMGLs was defined and depicted as a value of *p < 0.05, **p < 0.01, ***p < 0.001, and ****p < 0.0001, ns=not significant and while comparing different treatments of each genotype was defined and depicted as a value of #p < 0.05, ##p < 0.01, ###p < 0.001, and ###p < 0.0001, ns=not significant

### 3.6 Loss of *SORL1* decreases lysosomal exocytosis (LE) in hMGLs

Accumulation of substrates in lysosomes with impaired degradative capacity can add to further lysosomal stress. Lysosomal exocytosis (LE) is an alternate pathway to release lysosomes and their contents from cells (Wang et al., 2018; Zhong et al., 2023). LE is a calcium-dependent process that involves fusion of lysosomes with the plasma membrane, exposing the luminal domain of LAMP1 on the cell surface resulting in release of lysosomal contents into the extracellular environment (Andrews, 2000; Blott & Griffiths, 2002). Immune cells use LE to remodel the extracellular environment and to release some pro-inflammatory cytokines (Buratta et al., 2020; Tancini et al., 2020). Murine microglia are known to utilize LE to maintain cellular homeostasis and respond to external stimuli by releasing lysosomal enzymes and cytokines outside the cell (Andrei et al., 2004; Dou et al., 2012; Eder, 2009). We tested whether the process of LE might be altered because of lysosome dysfunction observed in *SORL1* KO hMGLs. We induced LE with the ionophore calcimycin (Jaiswal et al., 2002) and labeled cells with an antibody specific to the luminal epitope of LAMP1(Andrews, 2017) to visualize cell surface localization of LAMP1 via flow cytometry and immunocytochemistry. Measurement of surface LAMP1 localization has been repeatedly used as a robust readout of LE (Haka et al., 2016; Lee et al., 2024; Verma et al., 2023; Zhong et al., 2023). Both WT and *SORL1* KO hMGLs showed an increase in cell-surface localization of LAMP1 upon calcimycin treatment, suggesting that hMGLs can undergo LE *in vitro*. However, *SORL1* KO hMGLs exhibited reduced surface LAMP1 localization as compared to WT in both stimulated and unstimulated conditions, as evidenced by immunocytochemistry (Figure 6A) and flow cytometry (Figure 6B) showing that loss of *SORL1* results in decreased LE in hMGLs. This decrease was not due to the reduced total protein expression of LAMP1 (Figure 6C). Because LE causes extracellular release of lysosomal enzymes, we measured extracellular lysosomal enzyme activity and total levels of extracellular lysosomal enzymes. We found reduced extracellular enzyme activity of HEXB, Cathepsin B and Cathepsin D (Figure 6D-F) and reduced protein levels of all three lysosomal enzymes in cell culture media (Figure S3 A-D) in *SORL1* KO hMGLs upon treatment with a lysosomal exocytosis activator, calcimycin. While the reduction in extracellular enzyme activity and protein is likely due to the altered trafficking of hydrolases, taken together along with the reduction in cell surface LAMP1 localization this strongly indicates that important components of LE are impaired in *SORL1* KO hMGLs. Because the extracellular release of lysosomal enzymes is a normal part of LE, often contributing to extracellular matrix remodeling, cell-cell communication (Buratta et al., 2020) and ability to digest large extracellular substrates (Haka et al., 2016) the general reduction in lysosomal enzyme activity observed in *SORL1* hMGLs may further confound the LE process. To further confirm our hypothesis, we used fluorometric detection to measure extracellular lysosomal enzyme activity of HEXB and Cathepsin D as a readout of LE in a second model of hMGLs(Dolan et al., 2023). *SORL1* KO hMGLs showed decreased extracellular enzyme activity of both enzymes (Figure S2, C-D), validating the idea that loss of *SORL1* impairs LE in hMGLs irrespective of the method used to generate microglia from hiPSCs. To test whether LE contributes to secretion of cytokines and to assess contribution of SORLA to this process, we first assessed levels of a cytokine known to be released by LE in immune cells, IL-1β (Andrei et al., 2004; Gardella et al., 2001; Monif et al., 2016) and a cytokine known to be modulated by SORLA, IL-6 (Larsen & Petersen, 2017), with and without treatment with the LE activator, calcimycin. We measured both extracellular and intracellular levels of both cytokines upon treatment with DMSO, calcimycin and pro-inflammatory stimuli, LPS and IFN-y, Our data showed significant increase in extracellular secretion and no change in intracellular protein expression of both cytokines, upon treatment with calcimycin (Figure 6G,J; Figure S6A,D). These data suggest that calcimycin increases secretion of cytokines by increasing LE rather than by activating microglia. Interestingly we saw a slight decrease in the levels of IL-6 released after calcimycin treatment in *SORL1* KO hMGLs (Figure 6F), suggesting that altered levels of LE in these cells may affect release of this cytokine. Calcimycin-related secretion of IL1-β levels were unaffected in *SORL1* KO hMGLs, although intracellular protein levels of IL1-β were reduced (Figure S6).

**Figure 6.**
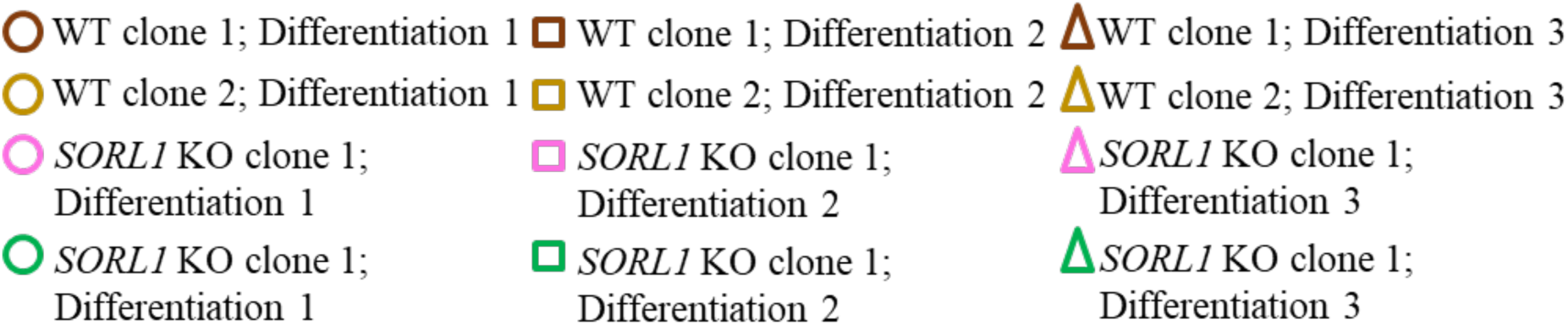
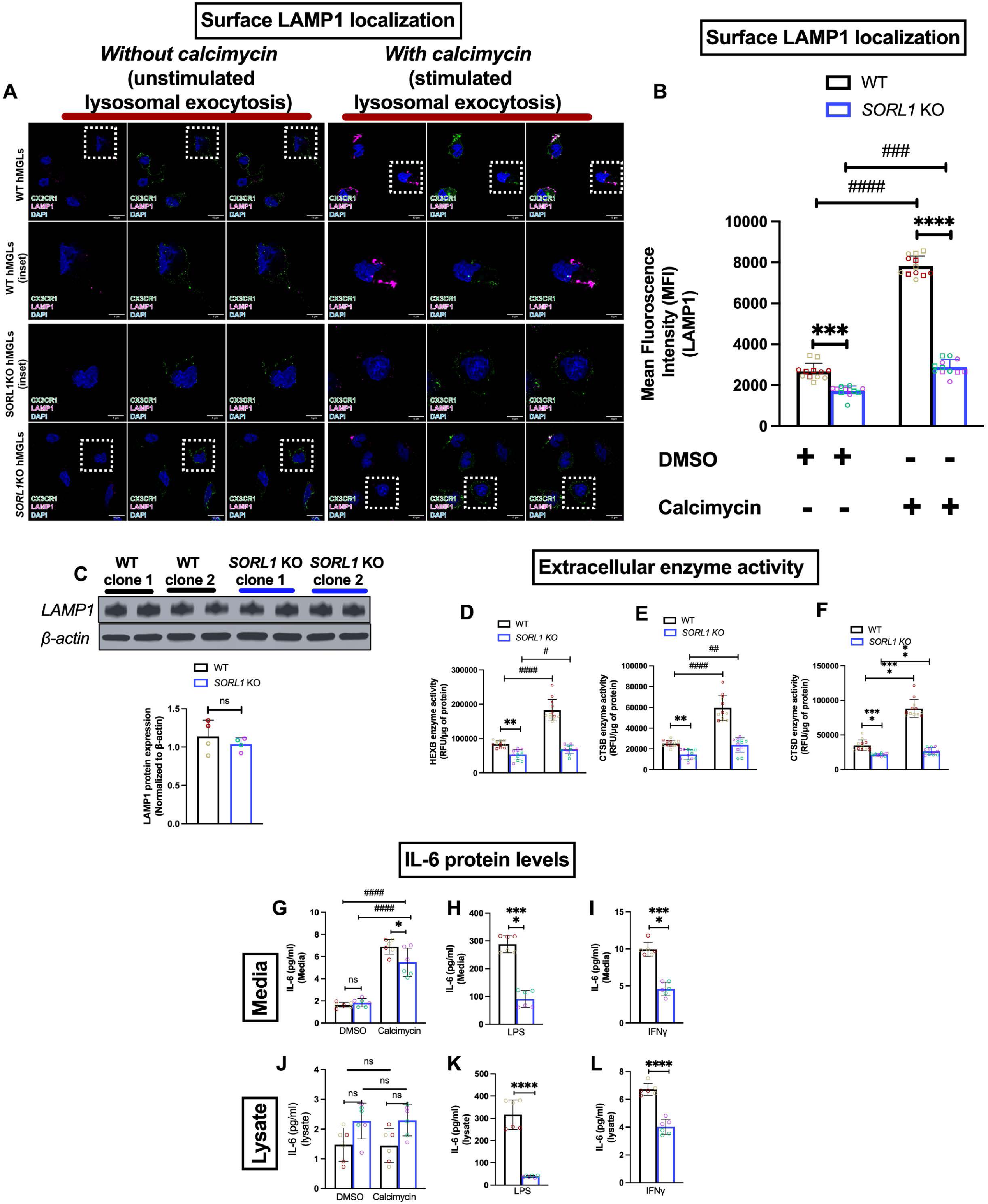
Loss of *SORL1* causes reduced lysosomal exocytosis in hMGLs. (A-B) Surface localization of LAMP1 measured by immunocytochemistry and flow cytometry as a readout of LE A) Immunocytochemistry with antibody specific for the N terminal luminal epitope of LAMP1 and microglia marker, CX3CR1 demonstrating decreased surface localization of LAMP1 in *SORL1* KO hMGLs as compared to WT hMGLs. B) Quantification by flow cytometry showing decreased MFI of LAMP1 antibody in *SORL1* KO hMGLs relative to WT hMGLs suggestive of decreased LE with loss of *SORL1* in hMGLs. C) Western blot and quantification shows no change in protein expression of LAMP1 in *SORL1*KO hMGLs relative to WT hMGLs. (D-F) Extracellular enzyme activity of lysosomal enzymes measured as a readout of LE. Enzyme activity assays demonstrate decreased lysosomal enzyme activity of D) HEXB, E) Cathepsin B and F) Cathepsin D in *SORL1* KO hMGLs as compared to WT hMGLs suggestive of decreased LE with loss of *SORL1* in hMGLs. (G-L) Cytokine secretion of IL-6 measured by ELISA shows increased extracellular secretion of IL-6 upon calcimycin treatment in both WT and *SORL1* KO hMGLs without increase in intracellular protein expression but *SORL1* KO hMGLs show decreased secretion of IL-6 in response to calcimycin as compared to WT hMGLs (G, J). *SORL1* KO hMGLs show reduced extracellular secretion and intracellular protein levels of IL-6 in response to pro- inflammatory stimuli LPS and IFNγ relative to WT hMGLs (H, I K, L). 2 isogenic clones per genotype (WT and *SORL1* KO) and 6 independent replicates (2 differentiations and 3 technical replicates) per clone per genotype (N=12 independent replicates) were used for these experiments. Data represented as mean ± SD and analyzed using parametric two-tailed unpaired t test and 2- way ANOVA with Tukeys multiple comparison test. Significance while comparing WT to *SORL1* KO hMGLs was defined and depicted as a value of *p < 0.05, **p < 0.01, ***p < 0.001, and ****p < 0.0001, ns=not significant and while comparing different treatments of each genotype was defined and depicted as a value of #p < 0.05, ##p < 0.01, ###p < 0.001, and ###p < 0.0001, ns=not significant.

### 3.6 Loss of *SORL1* results in overall decreased inflammatory response in hMGLs

While we noticed a slight but significant blunting of IL-6 in response to calcimycin treatment, we also observed a larger decrease in secretion and expression of IL-6 in response to canonical stimuli, LPS and IFNγ (Figure 6H,I,K,L). A recent study demonstrated that SORLA has a key role in sorting of the pattern recognition receptor CD14 (Ovesen et al., 2024) Similar to their findings, we also observed a blunted response of multiple pro and anti-inflammatory cytokines to LPS treatment and we also show a blunted response to IFNγ stimulation. Specifically, *SORL1* KO hMGLs showed decreased extracellular secretion of pro-inflammatory cytokines IL-1β, IL-15, IP-10, TNF-α and IL-8 and anti-inflammatory cytokines IL-10, IL-13, and IL-2 (Figure S6 and Figure S7). As aberrant lysosome function can negatively impact the neuroimmune response of microglia (J. D. Quick et al., 2023; Van Acker et al., 2021), our data along with that of Ovesen et al., suggests that there are several mechanism by which SORLA-related trafficking can affect pro-inflammatory responses that are important in AD.

## 4. Discussion

Despite extensive evidence that loss of function variants in the *SORL1* gene and *SORL1* haploinsufficiency are causative for AD (Campion et al., 2019; Holstege et al., 2017; Nicolas et al., 2016; Pottier et al., 2012; Verheijen et al., 2016), developing therapeutic interventions targeting *SORL1* has been challenging, partly because cell type-specific mechanisms through which loss of *SORL1* may increase risk for AD remain unclear. Indeed, recent RNA sequencing data highlights that certain CNS cells undergo more changes than others in response to *SORL1* KO (Lee et al., 2023). Recent work from our group and others has shown that depletion of SORLA in neurons leads to enlarged early endosomes, impairments in endo-lysosomal trafficking, changes in gene expression and impaired neuronal function (Hung et al., 2021; Knupp et al., 2020; Lee et al., 2023; Mishra et al., 2023; Mishra et al., 2022). Despite this extensive characterization in neurons, very little is currently understood about SORLA function in human microglia. Our previous work suggests that there may be important cell-type specific differences in endo-lysosomal organelles affected by loss of SORLA. We did not observe differences in the size early endosomes between SORL1 WT and SORL1 KO hMGLs (Knupp et al., 2020) but have documented lysosomal enlargement in SORL1 KO microglia (Mishra et al., 2024). Here we provide evidence for the role of SORLA in regulating lysosome function in human microglia. We show that loss of SORLA causes mis-trafficking of lysosomal enzymes, leading to impaired lysosomal degradation and accumulation of substrates in lysosomes. Alternate pathways to release lysosomal cargo from cells, such as lysosomal exocytosis, are also impaired. This impairment contributes to a blunted neuroimmune response.

SORLA binds to retromer (Fjorback et al., 2012), which regulates the retrograde transport of the cation independent mannose-6-phosphate receptor (CIMPR) through the Mannose-6-Phosphate dependent pathway (M6P) (Arighi et al., 2004; Cui et al., 2019). The CIMPR is a key receptor that traffics lysosomal enzymes from the TGN to the lysosomes (Thomas Braulke & Juan S Bonifacino, 2009; Braulke et al., 2023). In the absence of SORLA, there is reduced expression and reduced localization of CIMPR in late endosomes and lysosomes, suggesting that this crucial pathway for delivery of lysosomal hydrolases is impaired in *SORL1* KO hMGLs. During the process of lysosomal enzyme trafficking to the late endosomes and lysosomes, CIMPR forms a complex with premature lysosomal enzymes at the TGN, is transported to the early endosomes, then directed towards the lysosomes (Hasangaci et al 2015). Given that we do not see any difference in localization of this receptor at the TGN but reduced localization at the late endosomes and lysosomes, it is conceivable, that CIMPR gets stuck in the early endosomes during lysosomal enzyme trafficking in the absence of SORLA. In support of this idea, it has been shown in immortalized neuroblastoma and epithelial cell lines that SORLA can bind to the cytosolic adaptor protein, phosphofurin acidic cluster sorting protein 1 (PACS1)(Burgert et al., 2013) which is known to direct retrograde TGN retrieval of CIMPR(Chen et al., 1997; Scott et al., 2006; Wan et al., 1998) and can participate in PACS1 dependent sorting of Cathepsin B(Burgert et al., 2013; Nielsen et al., 2007). However, whether SORLA directly binds to CIMPR in and the precise role of SORLA in the M6P-dependent pathway in microglia warrants further investigation.

We did not observe significant differences in total protein or changes in pro vs. mature forms of lysosomal enzymes (Figure 2D-F). One explanation for this observation could be that there are different ratios of pro vs. mature forms of these enzymes in other compartments (such as endosomes) or that there is leakage of mature enzymes from the lysosome to the cytoplasm. Interestingly, proteomic analyses have variability in expression of lysosomal enzymes from *SORL1-*deficient microglia, in one case showing no change (Lee et al., 2023) and in another showing decreased expression although the majority of this decrease was reported from a cell line with a missense variant in the VPS10 domain of *SORL1* (Liu et al., 2020).

Impaired degradation in SORLA deficient hMGLs affects the fate of phagocytosed substrates. We observed increased internalization of various substrates including transferrin, fAβ and synaptosomes (Figure S5). Interestingly, we did not see a change in phagocytosis of oAβ, which replicates recently published work (Ovesen et al., 2024). The impact of increased phagocytosis of synaptosomes could contribute to excessive synaptic pruning and eventual synaptic loss observed in AD, but increased uptake of fAβ may be a means to reduce amyloidogenic burden. SORLA is known to regulate receptor protein homoeostasis at the cell surface by modulating intracellular sorting of receptors to the TGN and endo-lysosomal compartments (Pietilä et al., 2019; Schmidt et al., 2016). We observed increased cell surface localization of the phagocytic receptors P2Y6 and TREM2 (Figure S5) which may explain the increased internalization of certain substrates. Another recent study observed increased expression of the AD risk gene *BIN1* in *SORL1* KO hMGLs (Lee et al., 2023), which may also lead to increased phagocytosis (Sudwarts et al., 2022). Liu et al. observed expression changes in phagocytic receptors in SORL1 deficient hMGLs including *TREM2* and *ITGB2*, a receptor involved in the phagocytosis of both Aβ and synaptosomes (Liu et al., 2020). However, in their study, Liu et al. observed deficient phagocytosis of oligomeric Aβ, although the assay did not distinguish between phagocytic uptake and substrate degradation. Mechanisms of phagocytic uptake by microglia can also differ significantly depending on the protein conformation of Aβ(Pan et al., 2011; Ries & Sastre, 2016). (Lee et al., 2023; Sudwarts et al., 2022). Together, this data suggests that loss of *SORL1* expression may change localization of phagocytic receptors and genes involved in phagocytosis of specific substrates.

While we observe increased phagocytic uptake of both fAβ and synaptosomes, these substrates were not degraded, therefore a crucial aspect of the normal phagocytic process in *SORL1* KO hMGLs is impaired, leading to their intracellular accumulation. This may ultimately increase lysosomal stress in microglia and contribute to lysosomal dysfunction seen in AD. Indeed, the role of SORLA in microglial phagocytosis may act as a ‘double-edged sword’, functioning in both a disease-stage specific and context-dependent manner.

Lysosomal exocytosis (LE) is used by microglia to release cytokines, enzymes, and cargo into the extracellular space (Dou et al., 2012; Jacquet et al., 2024; Kreher et al., 2021) LE is a calcium-regulated process that expels lysosomal contents into the extracellular space (Buratta et al., 2020; Tancini et al., 2020). LE is important for such activities as remodeling of the extracellular matrix and, in the case of immune cells, releasing contents to degrade pathogenic material (Ireton et al., 2018; Miao et al., 2015). We found that *SORL1* KO hMGLs show reduced LE as evidenced by reduced LAMP1 localization on the plasma membrane and reduced enzyme activity in the conditioned medium of hMGLs after treatment with a stimulus. Thus, the increase accumulation of substrates in lysosomes may also indicate a failure to release these substrates via LE.

Interestingly, in response to an acute stress, LE can be used to release pro-inflammatory cytokines bypassing the slower, more conventional secretory pathway (Aiello et al., 2020). We observed that calcimycin-induced LE can release IL-1β and IL-6 with the IL-6 response being reduced in SORL1 KO hMGLs. We also observed a blunted cytokine release in response to canonical pro-inflammatory stimuli LPS and IFNγ, consistent with other recent reports in SORLA deficient hMGLs (Ovesen et al., 2024), suggesting that multiple mechanisms may govern SORLA-mediated cytokine release in hMGLs.

### 4.4 Working model

Altogether, we propose a working model that underpins the abnormal lysosomal phenotypes observed in SORL1 KO hMGLs (Figure 7). Loss of SORL1 impairs trafficking of lysosomal enzymes from the TGN to the lysosome via the M6P pathway, leading to inefficient degradation of phagocytosed substrates that accumulate in lysosomes. Alternate pathways of lysosomal clearance, such as LE, are also impaired. Furthermore, depletion of *SORL1* may decrease release of pro-inflammatory cytokines either through conventional secretory routes from the TGN to the cell surface or by impairing LE, as observed in our study and others (Ovesen et al., 2024). In the context of AD, we posit that lysosome dysfunction in microglia caused by loss of *SORL1* may result in heightened lysosomal stress caused by increased accumulation of substrates and stunted inflammatory response to accumulating pathogenic proteins in the extracellular brain environment. Since loss of *SORL1* is observed in early stages of AD, hypofunctional microglia with depleted SORLA may initially contribute to AD pathogenesis through lysosome dysfunction.

**Figure 7.**
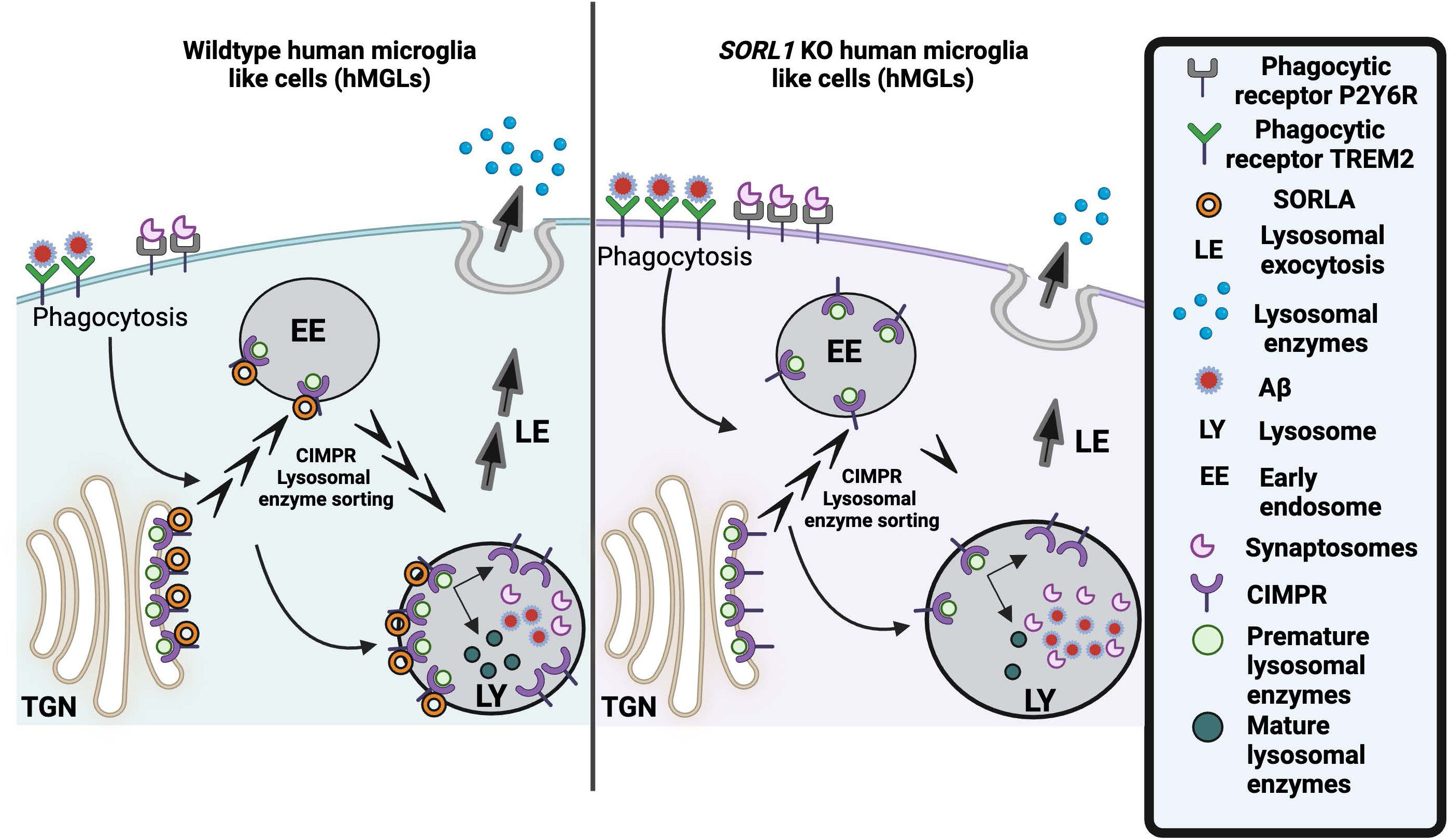
Working model. We propose the following working model that underpins the abnormal lysosomal phenotypes observed in *SORL1* KO hMGLs. Loss of SORL1 (orange circle) leads to decreased lysosomal degradation and decreased lysosomal enzyme activity caused by impaired trafficking of CIMPR and lysosomal enzymes from the TGN. Loss of SORL1 also causes enhanced phagocytic uptake due to increased cell surface localization of phagocytic receptors, but due to the decreased degradative capacity of the lysosomes, there is lysosomal accumulation of substrates including fibrillar Aβ 1-42 and synaptosomes. Alternate pathways of lysosomal clearance such as lysosomal exocytosis are also impaired, contributing to a blunted neuroimmune response. Altogether, loss of *SORL1* causes increased lysosomal stress in human microglia.

## Conclusions

In summary, our study highlights lysosome dysfunction in microglia as a pathway through which *SORL1* loss of function may increase the risk for AD. Our data identifies a previously unexplored cell type-specific mechanism through which *SORL1* can regulate the endo-lysosomal pathway beyond its well-established function as a sorting receptor for APP in neurons, illustrating the importance of exploring this multifaceted sorting receptor and potent AD risk factor as a therapeutic target for AD.

## Limitations of the study

We recognize our study has several limitations. *In vitro* studies using hiPSC-derived microglia do not allow thorough representation of cellular complexity characteristics of the physiological brain environment. We acknowledge that since our hMGLs are cultured in isolation, in the absence of neurons and astrocytes, and in two-dimensional cultures, their inflammatory response and morphology may be altered (Baxter et al., 2021). Future work should include studies with hMGLs transplanted into rodent brains or organoid models. Moreover, understanding the interaction of *SORL1* with other AD risk factors in regulating microglia lysosome function will further capture the complexity of AD. Nevertheless, our hiPSC model system has enabled a detailed investigation and provided evidence of cell autonomous role of *SORL1* in regulating microglial lysosome function.

## Supplementary information

### Supplementary methods

#### Differentiation of microglia from hiPSCs using Dolan et al 2023 protocol

hMGLs were also generated using a different iPSC-microglia differentiation protocol (Dolan et al., 2023). Briefly, hiPSCs were cultured in E8 media (Gibco #A1517001), dissociated with accutase and seeded in E8 media with rock inhibitor (RI; SelleckChem #S1049) at a density of 300k cells / well in ultra-low attachment plates (VWR #29443-030) to form embryoid bodies (EBs; DIV-1). EBs were cultured DIV0 to DIV4 in hypoxia (5% O2, 5% CO2, 37°C) and DIV4-DIV in normoxia (20% O2, 5% CO2, 37°C). EBs were given a full media change on DIV0 and cultured to DIV10 in iHPC media (1:1 IMDM:F12, 2% ITSG-X, 1% Glutamax, 1% Chemically-defined lipid concentrate, 1% NEAA, 10 ug/mL Poly(vinyl alchohol), 400 uM Monothioglycerol, 64 ug/mL L-ascorbic acid 2-Phosphate). On DIV0, EBs were treated with 50 ng/mL FGF2 (Gibco #PHG0261), 50 ng/mL BMP4 (Gibco #PHC9534), 12.5 ng/mL Activin-A (Gibco #PHC9564), 1 uM RI, 2 uM LiCl (Sigma #L7026). On DIV2, EBs were treated with 50 ng/mL VEGF (Peprotech #100-20), 50 ng/mL FGF2. On DIV4, DIV6, DIV8, EBs were treated with 50 ng/mL FGF2, 50 ng/mL VEGF, 10 ng/mL SCF (Gibco #PHC2116), 50 ng/mL TPO (Peprotech #300-18), 10 ng/mL IL-3 (Peprotech #200-03), 50 ng/mL IL-6 (Peprotech #AF-200-06). EBs were given a full media change on DIV0, DIV2, and DIV4 (2 mL / 6w), and media was added on top on DIV6 and DIV8 (1 mL / 6w). Floating iHPCs were separated from gravity-settled EBs by filtering through 40 um filter. iHPCs were counted with a Countess 3 (Invitrogen #AMQAX2000) and frozen in CryoStor CS10 (Stem Cell Technologies #07930). To differentiate into hMGLs, HPCs were thawed onto growth factor reduced matrigel-coated plates in 2 mL / 6w at a density of 100k / 6w in iMBM media (DMEM/F12, 2% ITS-G, 2% B-27, 1% N-2, 200 uM Monothioglycerol, 1% Glutamax, 1% NEAA, 5 ug / mL Insulin) with 50 ng/mL TGFB1 (Peprotech #100-21), 100 ng/mL IL-34 (BioLegend #577906), 25 ng/mL M-CSF (Peprotech #AF-300-25). hMGLs were given media additions with fresh cytokines (1 mL / 6w) every 2 days. hMGLs were re-plated on DIV22 and DIV30 at a density of 130k / 6w in 1:1 conditioned media and fresh media. hMGLs were re-plated by collecting floating cells, incubating attached cells in PBS for 5 minutes, and lifting attached wells with a cell lifter (VWR #29442-200). iMGL media was supplemented with 100 ng/mL CD200 (Bon Opus #C311) and 100 ng/mL CX3CL1 (Peprotech #300-31) beginning on DIV30 to promote microglial maturation.

#### Characterization of hiPSC-derived microglia generated using Dolan et al 2023 protocol

Differentiation of hMGLs from hiPSCs and HPCs was assessed at DIV22 (iMGL expression of CD33, P2RY12, CX3CR1, and high CD45) by flow cytometry. Surface markers were assessed by lifting cells and suspending in 5% BSA in PBS with 1:100 Fc Block (BioLegend Cat#422302). Cells were stained with pre-conjugated primary antibodies at 1:100 dilution for 15 min at 4°C.

Primaries antibodies used were: P2RY12-PE (BioLegend Cat# 392103), CX3CR1-PE (BioLegend Cat# 341603), CD33-PE/Cy7 (BioLegend Cat# 303402), and CD45-488 (BioLegend Cat# 304019). Cells were washed once with PBS and resuspended in 0.1% BSA in PBS with 1 μg/mL DAPI (Cayman Chemicals Cat# 14285-5). Flow cytometry data was collected on a Cytoflex LX analyzer (Beckman Coulter). Cells were identified by gating on FSC-A vs. SSC-A, singlets by FSC-A vs. FSC-H, and live cells by V450-A vs. SSC-A. Marker percentage positive was quantified by gating PE/Cy7 fluorescence channels using DAPI-only staining controls for each sample replicate.

#### Lysosomal enzyme activity assay using hiPSC-derived microglia generated using Dolan et al 2023 protocol

For enzyme activity assays performed with hMGLs generated using the protocol based on Dolan et al 2023, conditioned media was collected from hMGLs at DIV40, centrifuged at 300g for 5 minutes, and frozen at -80°C. hMGLs were lifted as described above and centrifuged at 300g for 5 minutes, supernatant removed, and cell pellets lysed in CD Lysis buffer (300 uL / 12-well). Lysates were centrifuged at 20,000g for 15 min at 4°C to pellet debris and then frozen at -80°C. Conditioned media and lysates were thawed on ice and aliquoted into a 96-well (Corning #3904; 50 uL/well, 2 replicate wells/sample). CTSD substrate (Abcam #ab65300) or HEXB substrate (Cell Biolabs #MET-5095) was added to wells (50 uL/well) and plates were incubated at 37°C for 30 minutes. Neutralization reagent (100 uL/well) was added to HEXB assay wells. Sample fluorescence was measured using a Perkin Elmer EnSpire Plate Reader. CTSD: Ex 328 / Em 460; HEXB: Ex 365 / Em 460. Fluorescence values were averaged between replicate wells.

## Acknowledgments

We acknowledge members of the Young and Jayadev laboratories for helpful critiques and discussions of this work. We also acknowledge Dr. Anna Kane in the Stevens lab for critical reading and editing of the manuscript.

## Data Availability

The data that support the findings of this study are available from the corresponding author upon reasonable request.

## Author Contributions

Conceptualization: SM, JEY, SJ; Experimental Methodology: SM, NM; SS, CK. Writing, original draft: SM, JEY; Writing, reviewing and editing: SM, NM, BS, SJ, JEY; Funding acquisition: SM, NM, BS, JEY

## Supplementary Figure legends

**Supplementary figure 1.**
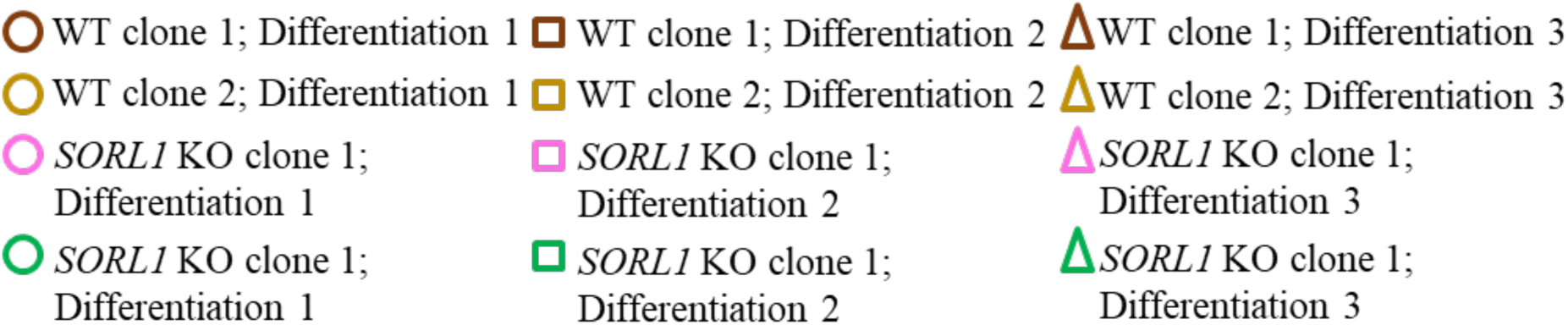
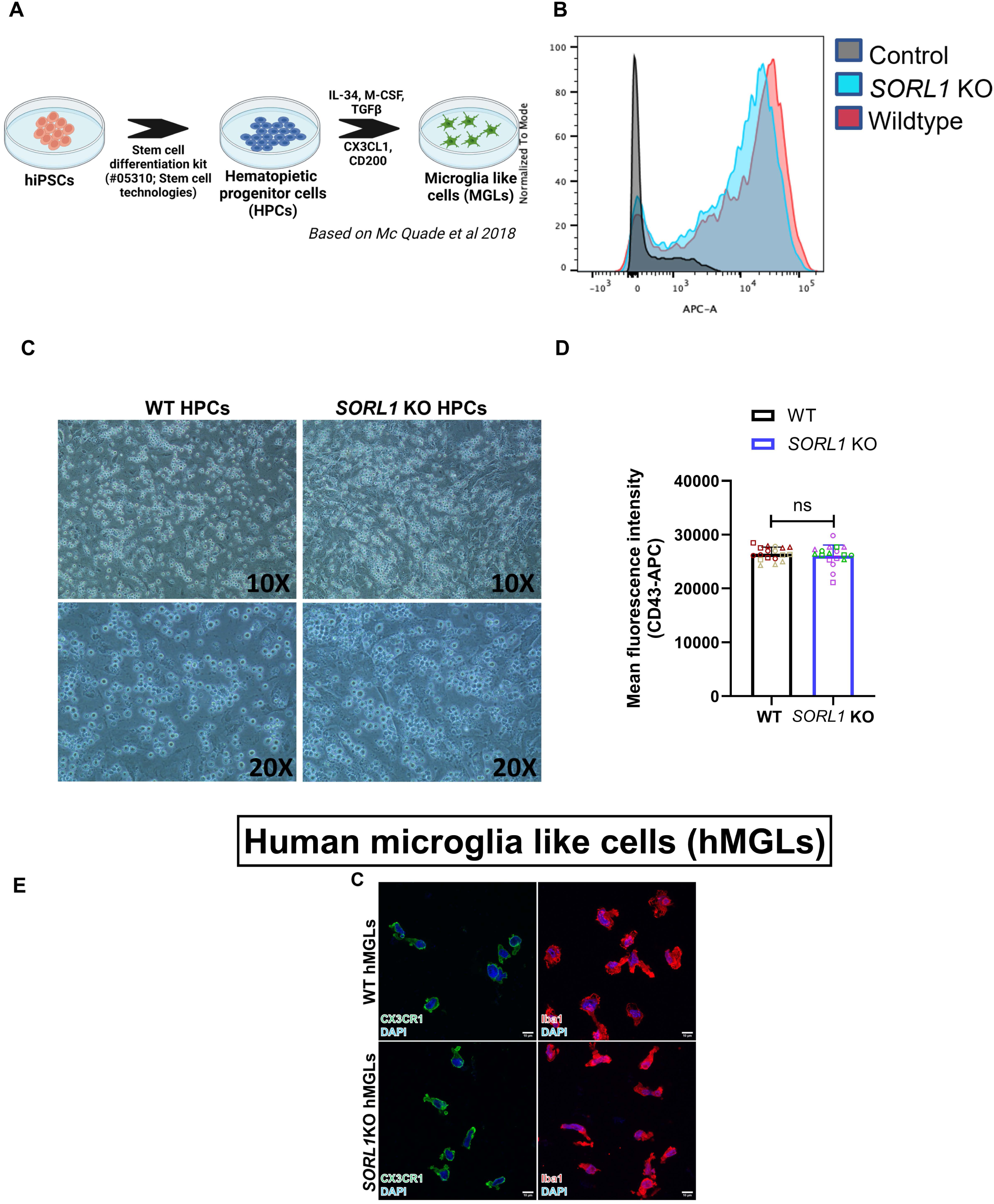
Characterization of Hematopoietic progenitor cells (HPCs) and microglia derived from hiPSCs (hMGLs) with loss of *SORL1* (A) Schematic showing protocol used for derivation of HPCs and microglia from hiPSCs. Protocol was based on (McQuade et al., 2018) with some modifications. (B) Histogram generated using FlowJo showing positive expression of HPC marker, CD43 in both WT and *SORL1* KO hMGLs. (C) Brightfield images at 10X and 20X magnifications, showing HPCs (bright grape like clusters) derived from WT and *SORL1* KO hiPSCs. (D) Mean fluorescence intensity measured by flow cytometry showing no difference in expression of HPC marker, CD43 in WT and *SORL1* KO hMGLs. (E) Immunocytochemistry showing positive expression of microglia specific markers including Iba1, and CX3CR1 in both WT and *SORL1* KO hMGLs. Images were acquired using confocal microscopy and Leica software. Scale bar = 10μm. 2 isogenic clones per genotype (WT and *SORL1* KO) and 9 independent replicates per clone per genotype (N=18 independent replicates) were used for these experiments. Data represented as mean ± SD and analyzed using parametric two-tailed unpaired *t* test. Significance was defined as a value of \**p* < 0.05, \*\**p* < 0.01, \*\*\**p* < 0.001, and \*\*\*\**p* < 0.0001, ns=not significant.

**Supplementary figure 2.**
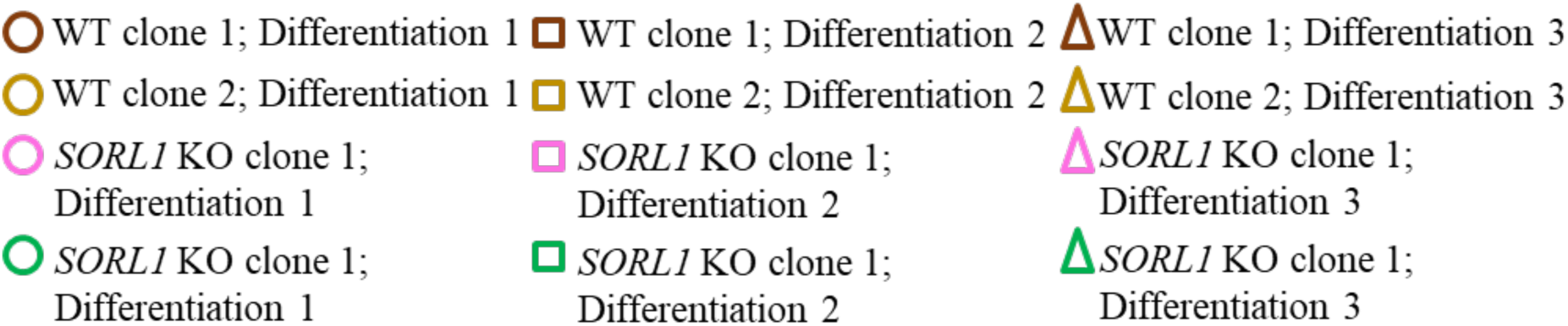
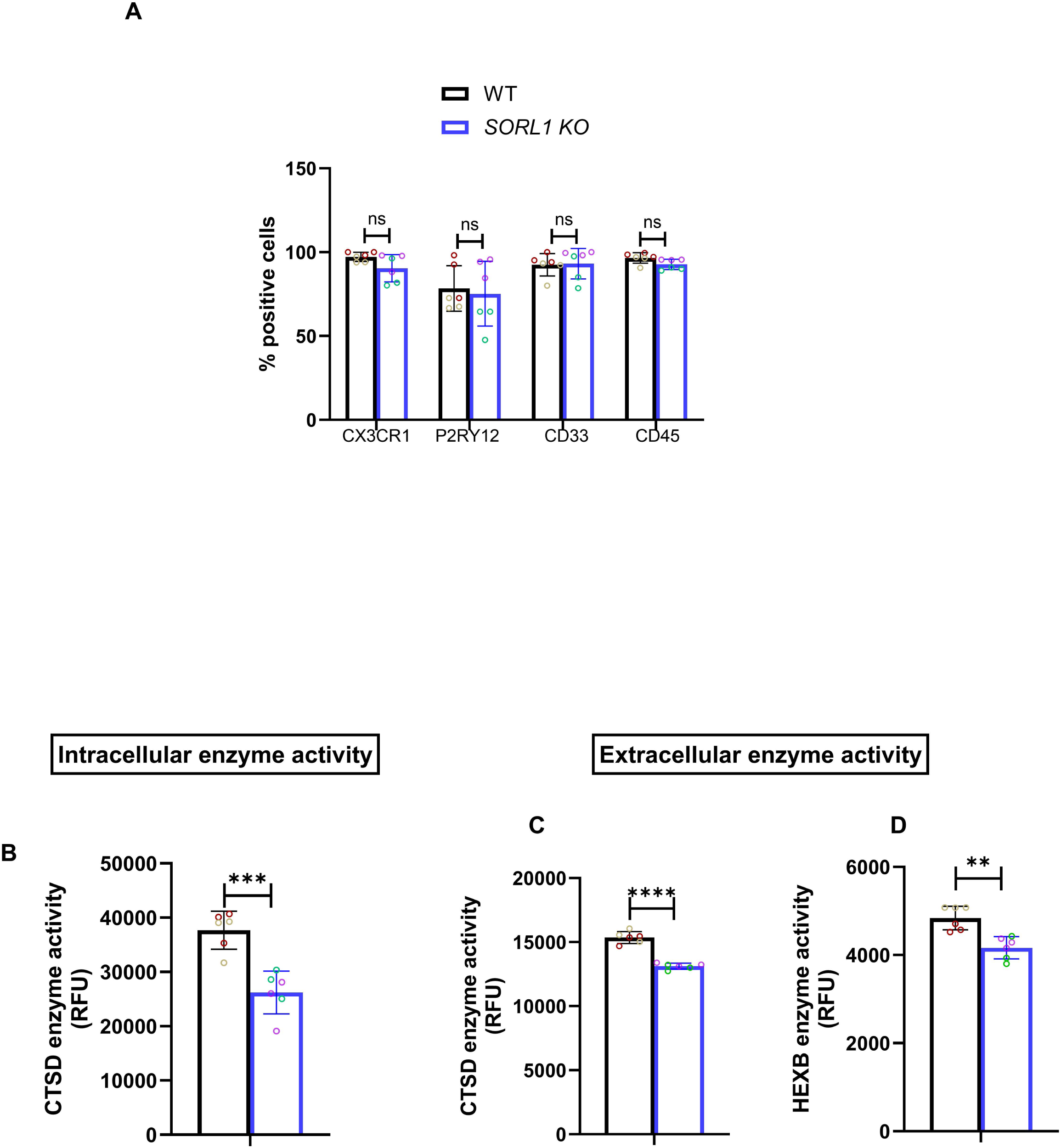
Loss of *SORL1* results in reduced intracellular enzyme activity of CTSD and decreased extracellular enzyme activity of CTSD and HEXB lysosomal enzymes in hMGLs generated using Dolan et al protocol. (A) Expression of microglia specific markers, CD33, P2RY12, CX3CR1 and CD45 high assessed by flow cytometry. Enzyme activity assays showing reduced intracellular enzyme activity of CTSD (B) and reduced extracellular enzyme activity of CTSD and HEXB (C-D) indicating reduced LE in *SORL1* KO hMGLs as compared to WT hMGLs. 1 clone per genotype (WT and *SORL1* KO) and 6 independent replicates per clone per genotype (N=6 independent replicates) were used for these experiments. Data represented as mean ± SD and analyzed using parametric two-tailed unpaired *t* test. Significance was defined as a value of \**p* < 0.05, \*\**p* < 0.01, \*\*\**p* < 0.001, and \*\*\*\**p* < 0.0001, ns=not significant.

**Supplementary figure 3.**
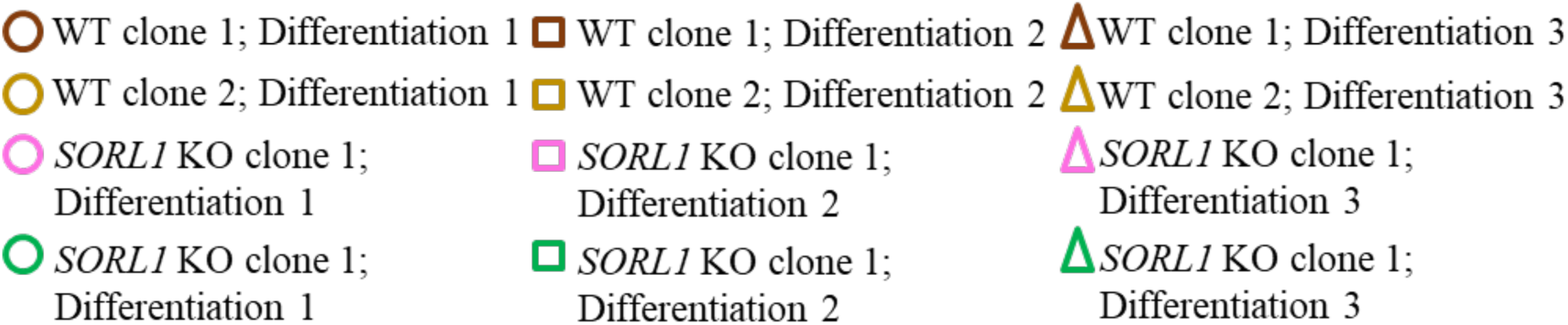
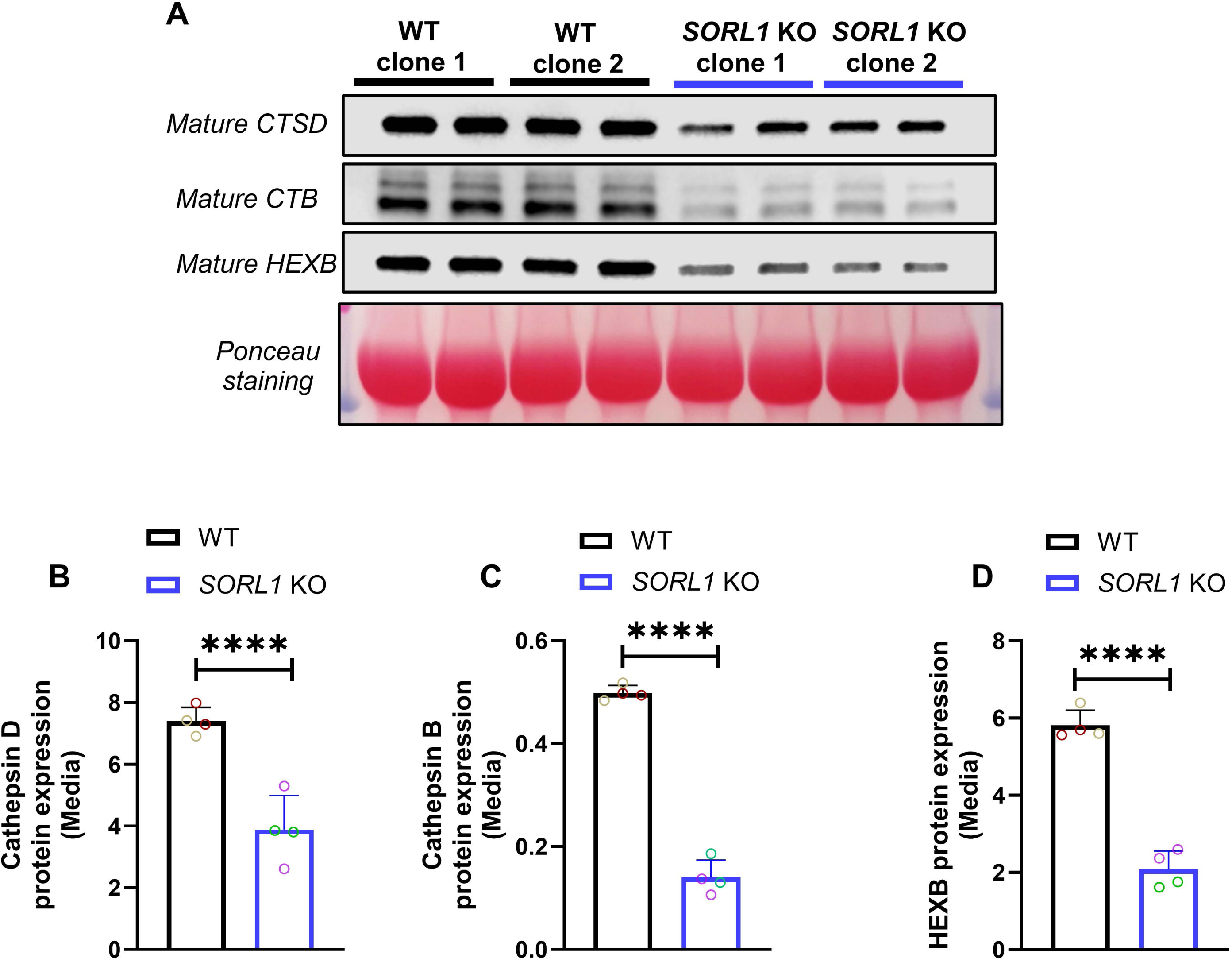
Loss of *SORL1* causes reduced extracellular release of lysosomal enzymes in hMGLs. (A) Western blotting showing decreased extracellular levels of the lysosomal enzymes, CTSD, HEXB and CTSB after treatment with the lysosomal exocytosis activator, calcimycin. (B) Quantification of western blotting data using Image J software. 2 isogenic clones per genotype (WT and *SORL1* KO) and 2 independent replicates per clone per genotype (N=4 independent replicates, 1 differentiation) were used for these experiments. Data represented as mean ± SD and analyzed using parametric two-tailed unpaired *t* test. Significance was defined as a value of \**p* < 0.05, \*\**p* < 0.01, \*\*\**p* < 0.001, and \*\*\*\**p* < 0.0001, ns=not significant.

**Supplementary figure 4.**
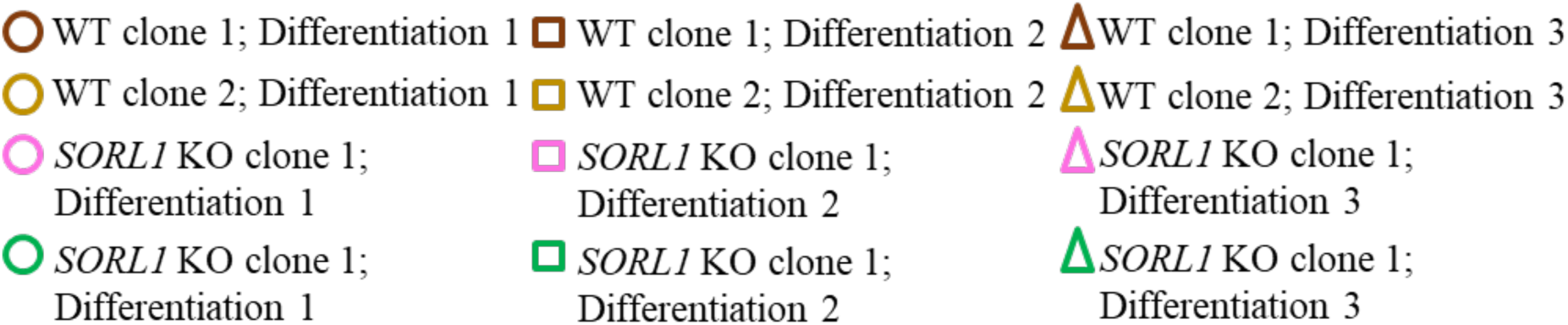
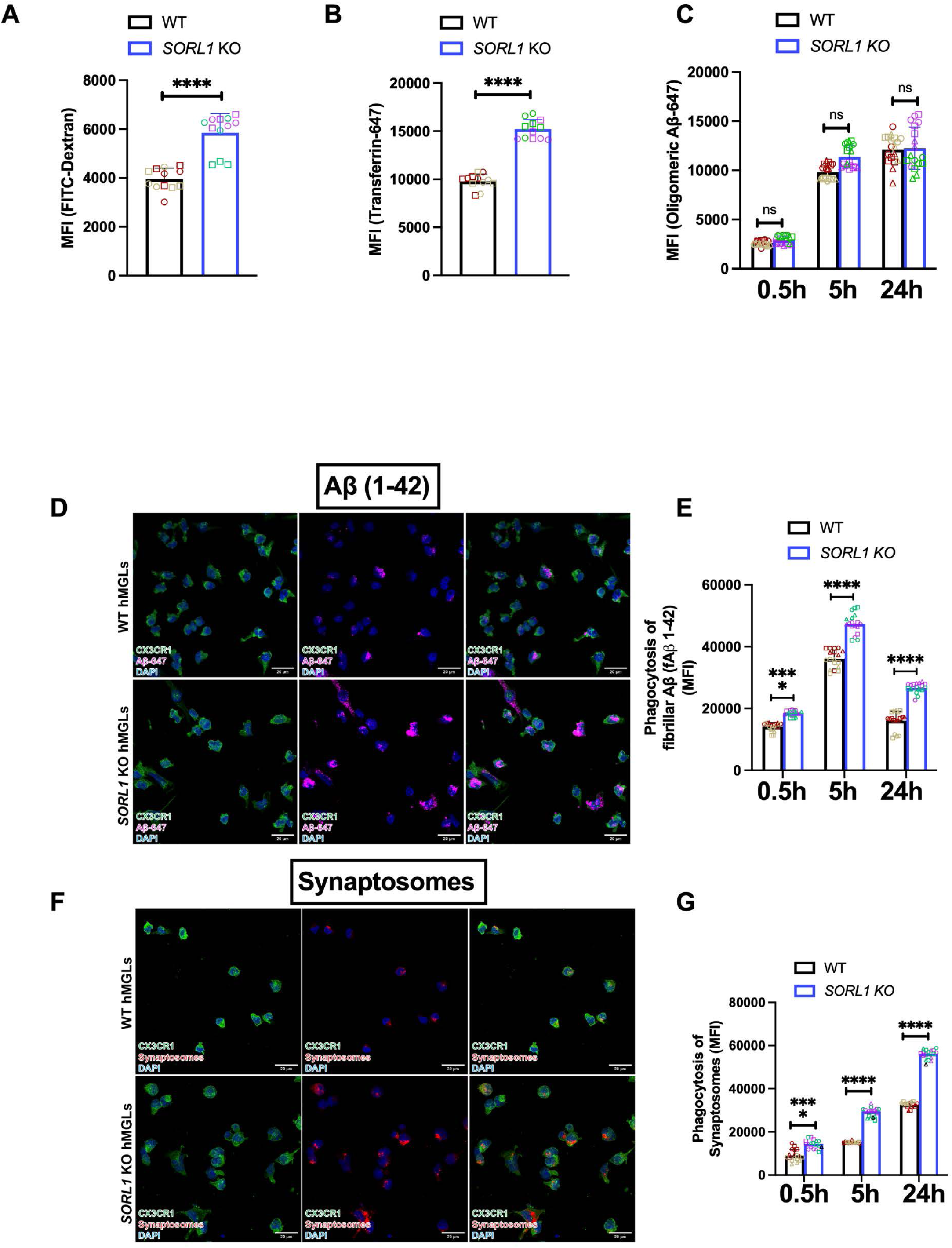
*SORL1* alters phagocytosis in a substrate specific manner. (A) Uptake assay of pinocytosis substrate, FITC-Dextran using flow cytometry showing increased intracellular fluorescence intensity of FITC-Dextran in *SORL1* KO hMGLs as compared to WT hMGLs. (B) Uptake assay of an endocytosis marker, transferrin conjugated with fluorophore, Alexa fluor 647, using flow cytometry showing increased intracellular fluorescence intensity of Transferrin-647 in *SORL1* KO hMGLs as compared to WT hMGLs. (C) Phagocytosis assay with Oligomeric Aβ-647 (oAβ) with cells treated for 30 mins, 5h and 24h and quantification of intracellular Mean fluorescence intensity (MFI) using flow cytometry, as a readout of phagocytosis in hMGLs demonstrating increased phagocytosis in SORL1 KO hMGLs relative to WT hMGLs. Immunocytochemistry with hMGLs treated with fibrillar Aβ-647 and CM-Dil dye labeled synaptosomes and co-labeled with microglia marker, CX3CR1 showing increased intracellular phagocytosis of (D) fibrillar Aβ-647 and (F) synaptosomes in *SORL1* KO hMGLs as compared to WT hMGLs. Quantification of Mean fluorescence intensity (MFI) as a readout of phagocytosis in hMGLs treated with both substrates for 30 mins, 5h and 24h demonstrating increased phagocytosis of (E) fibrillar Aβ-647 and (G) synaptosomes in *SORL1* KO hMGLs relative to WT hMGLs. Scale bar = 10μm. 2 isogenic clones per genotype (WT and *SORL1* KO) for all the experiments. 6 independent replicates (2 differentiations and 3 technical replicates) were used for the FITC-Dextran and Transferrin-647 uptake assays and 9 independent replicates per clone per genotype (N=18 independent replicates, 3 differentiations and 3 technical replicates) were used phagocytosis assays of fAβ and oAβ. Data represented as mean ± SD and analyzed using parametric two-tailed unpaired *t* test. Significance was defined as a value of \**p* < 0.05, \*\**p* < 0.01, \*\*\**p* < 0.001, and \*\*\*\**p* < 0.0001, ns=not significant.

**Supplementary figure 5.**
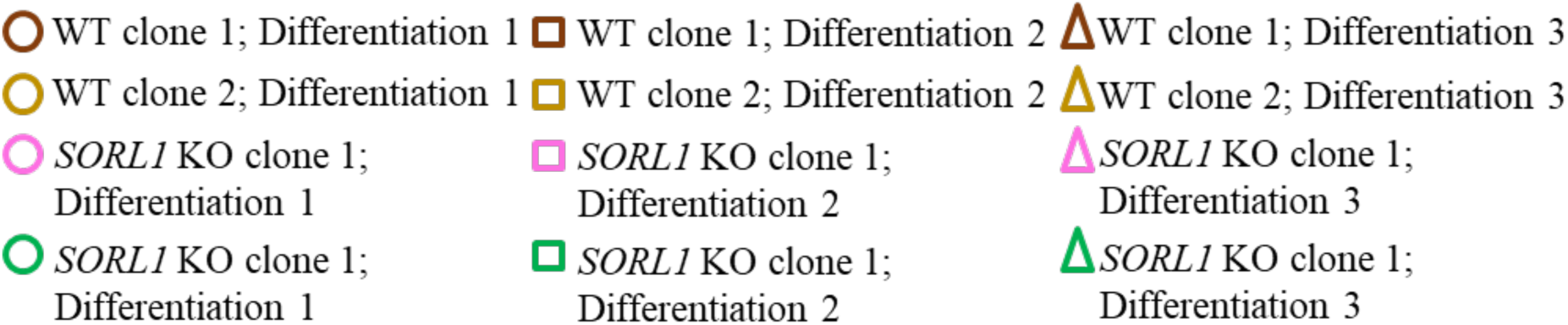
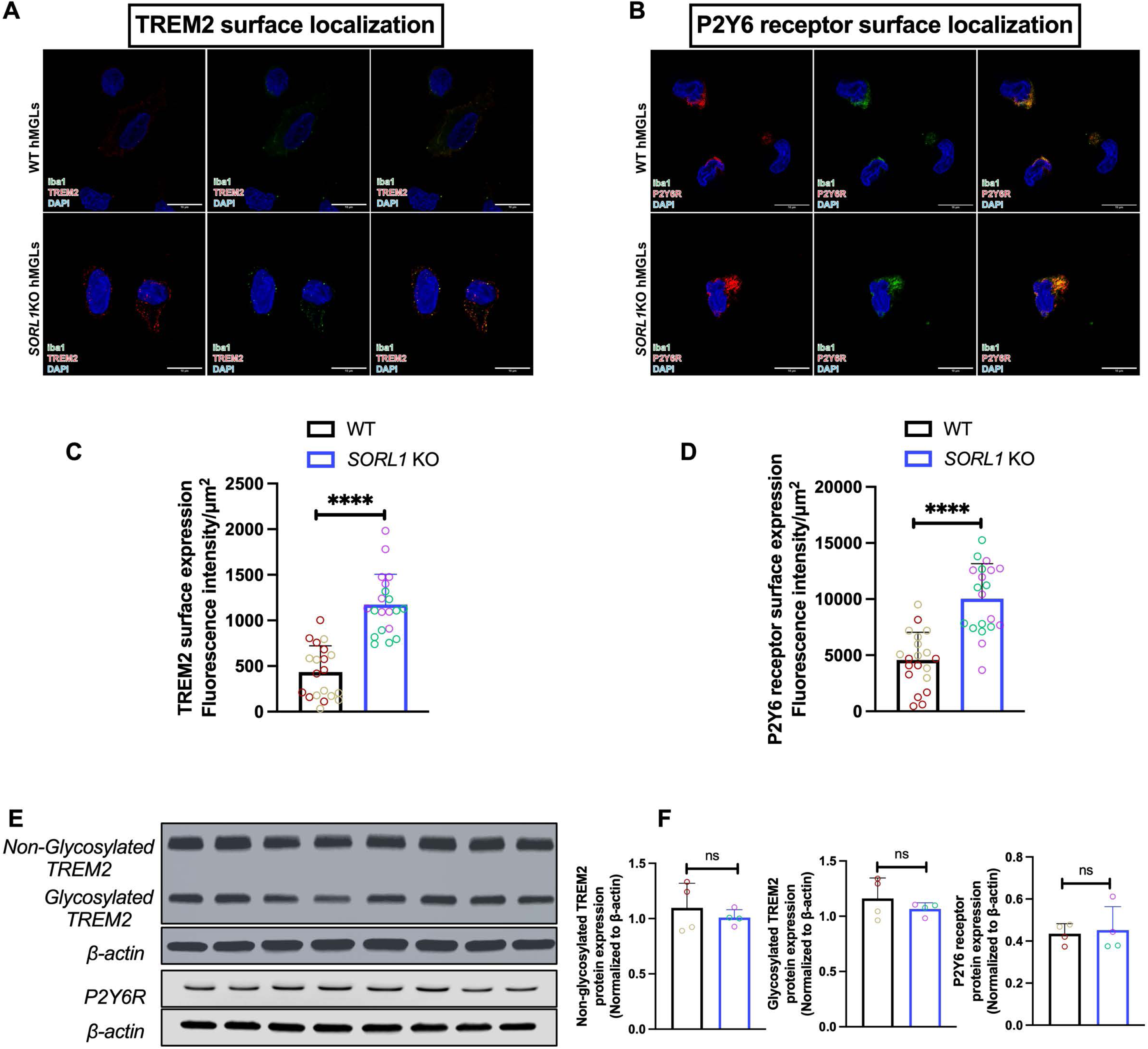
Loss of *SORL1* results in increased surface localization of phagocytic receptors TREM2 and P2Y6 receptor. Cell surface localization of phagocytic receptorsTREM2 (A,C) and P2Y6 (B,D) receptor measured by immunocytochemistry and Image J software showing increased surface localization of both phagocytic receptors in *SORL1* KO hMGLs as compared to WT hMGLs. Data represented as Mean fluorescence intensity per cell area in µm2. (E-F) Western blotting shows no change in TREM2 and P2Y6 receptor protein expression in *SORL1* KO hMGLs as compared to WT hMGLs. Images were acquired using confocal microscopy and Leica software. Scale bar = 10μm. 2 isogenic clones per genotype (WT and *SORL1* KO; 1 differentiation per clone per genotype) were used for these experiments. Data represented as mean ± SD and analyzed using parametric two-tailed unpaired *t* test. Significance was defined as a value of \**p* < 0.05, \*\**p* < 0.01, \*\*\**p* < 0.001, and \*\*\*\**p* < 0.0001, ns=not significant.

**Supplementary figure 6.**
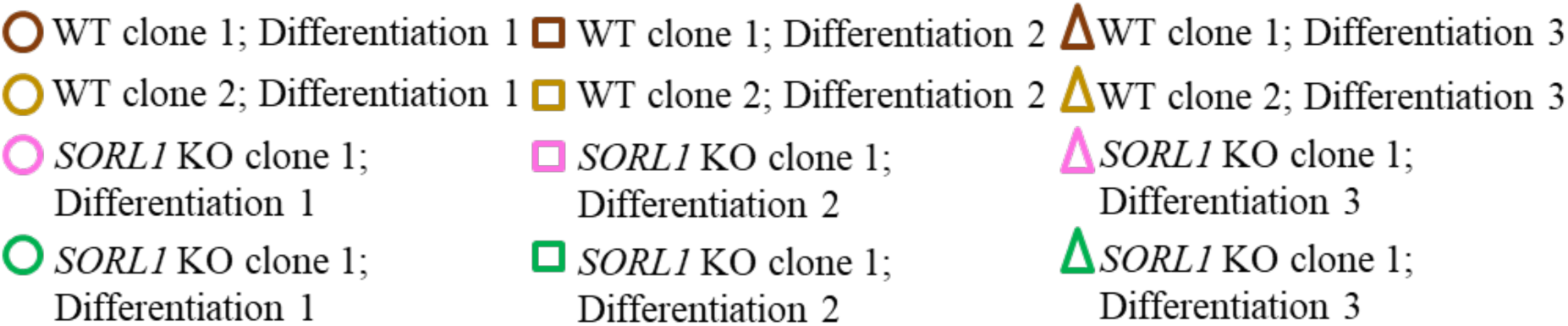
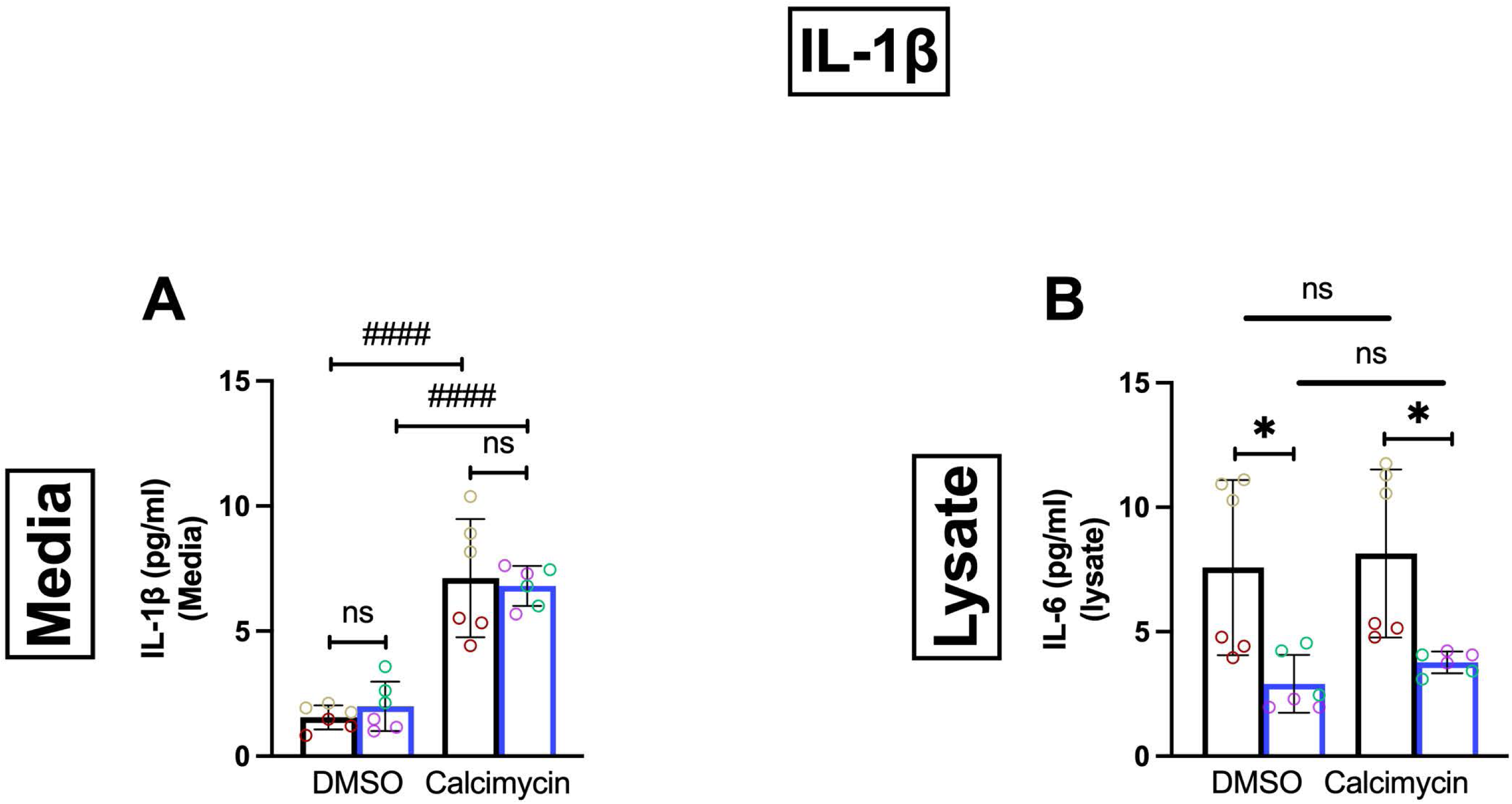
hMGLs secrete IL-1β through lysosomal exocytosis and *SORL1* deficiency does not change this secretion. (A-B) ELISAs measuring extracellular (media) and intracellular (lysate) IL-1β cytokine levels in cells treated with and without calcimycin showing increased secretion of IL-1β with calcimycin (LE activator). Calcimycin treatment did not alter levels of IL1-β in the cell lysate although *SORL1* KO hMGLs did show reduced IL1-β expression as compared to WT hMGLs. 2 isogenic clones per genotype (WT and *SORL1* KO; 1 differentiation per clone per genotype) were used for these experiments. Data represented as mean ± SD and analyzed using parametric two-tailed unpaired *t* test. Significance was defined as a value of \**p* < 0.05, \*\**p* < 0.01, \*\*\**p* < 0.001, and \*\*\*\**p* < 0.0001, ns=not significant.

**Supplementary figure 7.**
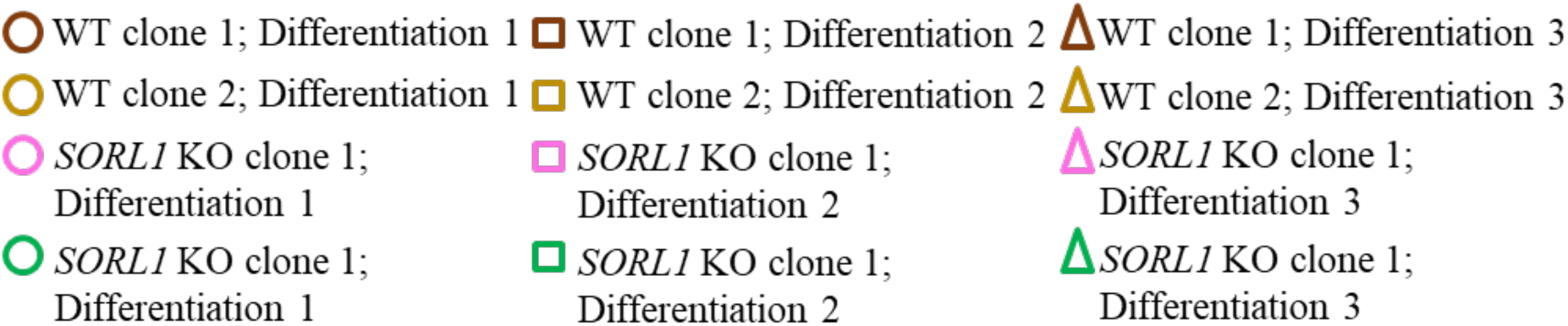
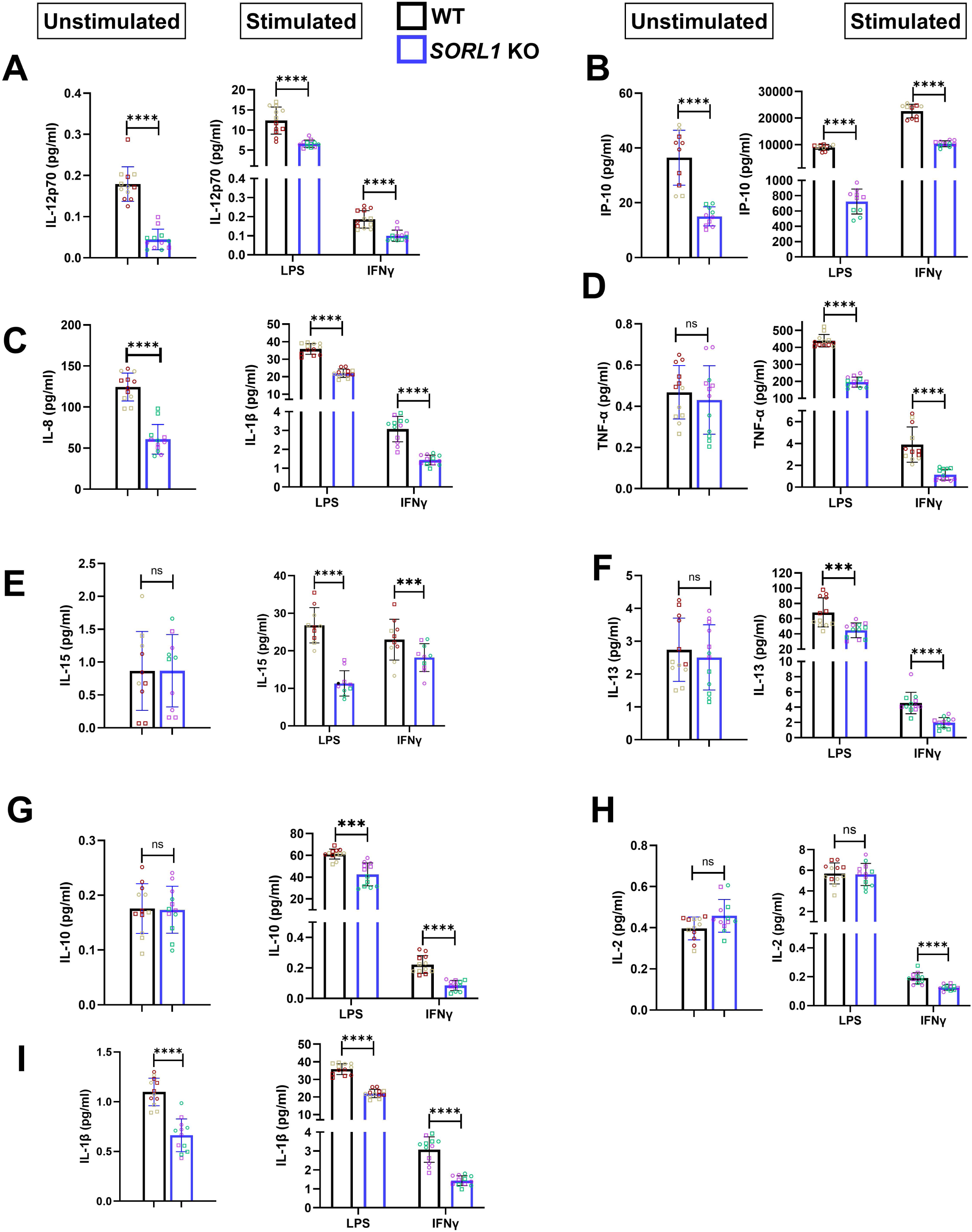
Loss of *SORL1* results in reduced inflammatory response to pro-inflammatory stimuli in hMGLs. Cytokine levels measured from the media of hMGS after treatment with pro-inflammatory stimuli LPS and IFN-γ. In general there is a blunted response in *SORL11* KO hMGLS. These cytokines include (A) IL-12p70; (B) IP-10; (C) IL-8; (D) TNFα, (E) IL-15; (F) IL-13; (G) IL-10; (H) IL-2; (I) IL-1β. Unstimulated *SORL1* KO hMGLs demonstrated reduced extracellular levels of (A-C) IL-2p70 (IL-12), IP-10, IL-8 and IL-1β but no change in (D-H) TNF-α, IL-15, IL-13, IL-10 and IL-2 as compared to WT hMGLs. 2 isogenic clones per genotype (WT and *SORL1* KO) and 6 independent replicates (2 differentiations and 3 technical replicates) per clone per genotype (N=12 independent replicates) were used for these experiments. Data represented as mean ± SD and analyzed using parametric two-tailed unpaired *t* test. Significance was defined as a value of \**p* < 0.05, \*\**p* < 0.01, \*\*\**p* < 0.001, and \*\*\*\**p* < 0.0001, ns=not significant.

